# Formation of an Enduring Ensemble of Accumbens Neurons Leads to Prepotent Seeking for Cocaine Over Natural Reward Cues

**DOI:** 10.1101/2024.08.05.606522

**Authors:** Reda M Chalhoub, Anze Testen, Jordan Hopkins, Camille Carthy, Peter W Kalivas

**Affiliations:** Department of Neuroscience, Medical University of South Carolina, Charleston, South Carolina, USA; Medical Scientist Training Program, Medical University of South Carolina, Charleston, South Carolina, USA; Ralph Johnson Veterans Administration, Charleston, South Carolina, USA

**Keywords:** Nucleus Accumbens, Reward encoding, Calcium Imaging, Cocaine, Medium Spiny Neurons, Ensembles

## Abstract

Neuronal activity in the nucleus accumbens core (NAcore) is necessary for reward-seeking behaviors. We hypothesized that the differential encoding of natural and drug rewards in the NAcore contributes to substance use disorder. We leveraged single-cell calcium imaging of dopamine D1- and D2-receptor-expressing medium spiny neurons (MSNs) in the NAcore of mice to examine differences between sucrose and cocaine rewarded (self-administration) and unrewarded (abstinent and cue-induced) seeking. Activity was time-locked to nose-poking for reward, clustered, and compared between sucrose and cocaine. Only in cocaine-trained mice were excited D1-MSNs securely stable, capable of decoding nose-poking in all rewarded and unrewarded sessions and correlated with the intensity of nose-poking for unrewarded seeking. Furthermore, D1-MSNs formed a stable ensemble predictive of seeking behavior after extended cocaine, but not sucrose abstinence. The excited D1-MSN ensemble uniquely drives cue-induced cocaine seeking and may contribute to why drug seeking is prepotent over natural reward seeking in cocaine use disorder.

## INTRODUCTION

A major goal of research into the neural mechanisms of substance use disorder (SUD) is to understand why a person suffering from SUD chooses to relapse to drug use at the expense of engaging in nondrug rewarding life experiences. We hypothesized that an idea put forth over 30 years ago that discrete neuronal subpopulations within the brain, termed ensembles, encode specific learned behaviors might offer insight into the underpinnings of this cardinal symptom of SUDs^1^. The ensemble hypothesis has been invigorated over the last decade by advances in visualizing and manipulating genetically discrete neuronal subgroups^2^. One approach uses transgenic rodents transfected with immediate early gene (IEG) promoters that can mark neuronal ensembles activated by initiating behaviors^3,4^. Also, advances in techniques to record the activity of single neurons in freely behaving animals, including in-vivo electrophysiology^5,6^ or single-cell resolution calcium (Ca^2+^) imaging^7,8^, opened avenues for ensemble characterization.

In the field of SUDs research, the IEG approach reveals that drug seeking initiated by drug-associated context and discrete cues requires activity in a relatively small ensemble of neurons (2-3%) in the nucleus accumbens^4,9,10^. The nucleus accumbens is a well-established nexus in processing both drug and natural reward seeking and is composed largely of two genetically distinct subpopulations of medium spiny neurons expressing either D1 (D1-MSN) or D2 dopamine receptors (D2-MSN)^11^. Importantly, the ensembles of IEG positive neurons induced by either cocaine- or sucrose-conditioned cues overlap by only ∼25%, indicating that the nucleus accumbens uses largely separate ensembles to facilitate cue-initiated drug versus natural reward seeking^4^, a finding consistent with earlier in vivo electrophysiological measurements^12^.

The IEG approach examines only neurons that are excited and does not consider neurons inhibited by a stimulus, nor does it quantify time-specific changes in individual cell activity across task performance^13^. For these reasons, we employed in vivo single cell Ca^2+^ imaging in male and female mice expressing Cre recombinase in either D1- or D2-MSNs. We compared neuronal ensembles created during cocaine or sucrose self-administration (rewarded seeking) and during two unrewarded seeking sessions; after a period of abstinence and in response to reward-associated cues after extinction training. This approach allowed us to record increases and decreases in activity of accumbens D1- and D2-MSNs that were time-locked to a behavioral operand for seeking behavior, active nose-pokes (NPs). Furthermore, we leveraged the spatial resolution of calcium imaging to investigate and contrast the stability of recorded neurons over the time-course of a single seeking session and longitudinally between seeking sessions.

## RESULTS

### Behavioral Responses

Mice were virally transfected with a Ca^2+^ reporter (pAAV-Syn-Flex-GCaMP6f-WPRE-SV40) into the core subcompartment of the nucleus accumbens (NAcore) and 4-8 weeks later implanted with a lens over the viral injection site (Figure 1a-c)^14^. Transfected neurons were visible in 26 of 35 mice, which were divided into two groups and trained for 12 days to self-administer either cocaine or sucrose pellets paired with a light/tone cue (Figures 1d-f). During sucrose or cocaine self-administration, active NPs were reinforced by a sucrose pellet or cocaine infusion when NPs were separated by a 20 sec time-out period, while inactive NPs never delivered a reward. Following 10 days of forced abstinence, mice were returned to the operant chamber for a session with cues but no reinforcer (termed post-abstinence seeking) to test for drug seeking, followed by 5-10 days of extinction training to criterion without cue or reinforcer presentation. After extinction training was completed, mice underwent cued reinstatement, during which active NPs by extinguished mice yielded cue only. Calcium recordings were made during the following sessions, late (stable) self-administration (two separate recordings across days 8-12), during the first day post-abstinence (PA) and during the final cue-only reinstatement session (Figure 1d). D1- and D2-Cre mice showed equivalent levels of self-administration across reward and genotype modalities (Figures 1e,f). Similarly, equivalent rewarded active nose-poking between D1- and D2-Cre mice was observed during the PA session and extinction training (Figures 1g,h and S1a,c). However, akin to previous findings^4,15^, D1- and D2-Cre mice reinstated more to cocaine cues than sucrose cues (Figure 1i). Inactive NPs were not different between D1- and D2-Cre mice in rewarded (Figure 1e,f) or unrewarded (Figure S1) seeking. No sex differences appeared in any of the behavioral measures shown in Figure 1 (Figure S2).

**Figure 1.**
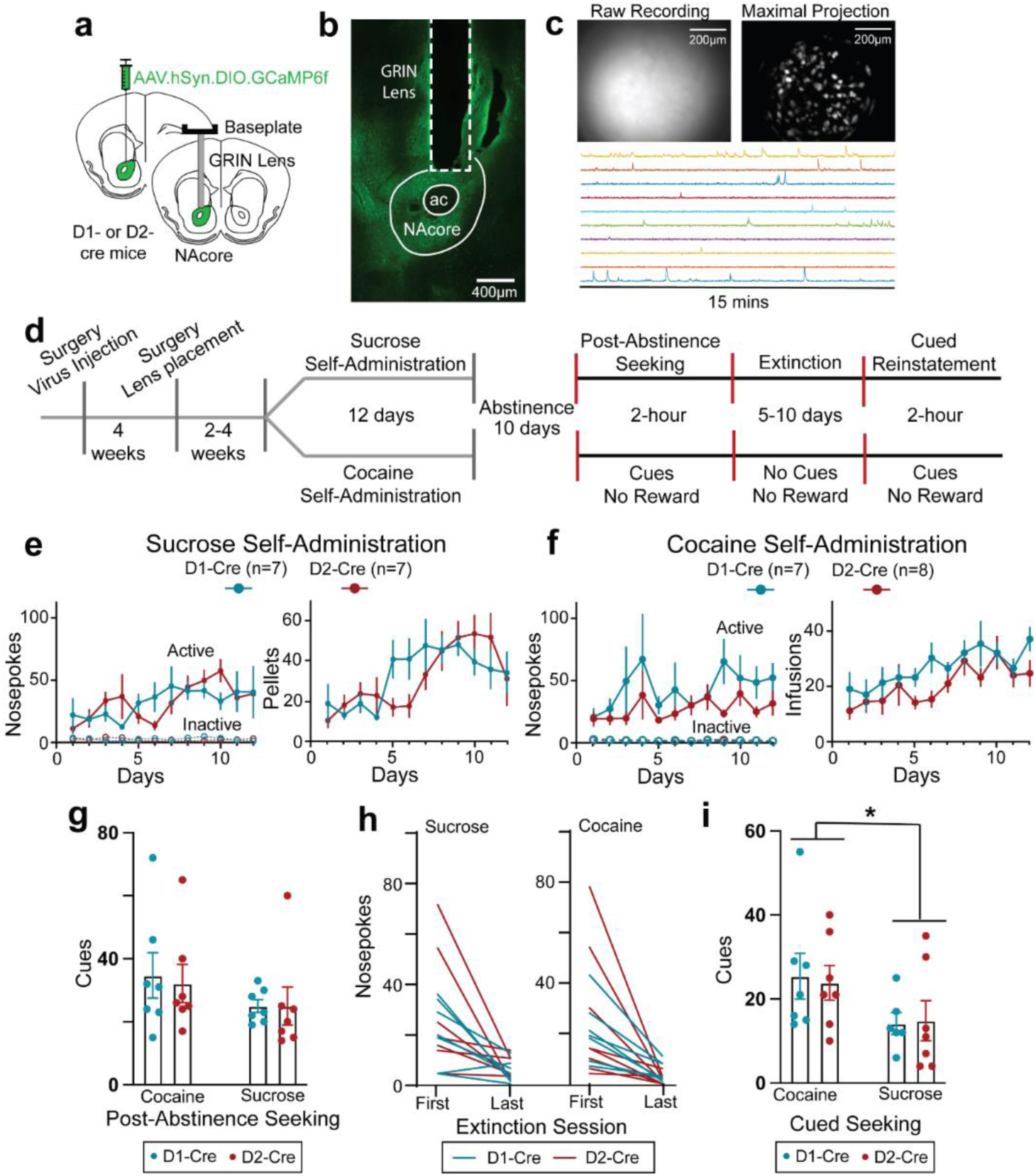
Single-cell resolution imaging of D1- and D2-MSN of nucleus accumbens core during sucrose/cocaine rewarded and unrewarded seeking. **(a)** Illustration showing the surgical planning, including virus injection location and planned lens placement in D1- and D2-cre mice. **(b)** Micrograph from D1-cre mouse showing virus expression (stained with GFP+, green) and gradient index (GRIN) lens tract. **(c)** Field of view recording of a D1-cre mouse (upper left panel), the maximal intensity projection of the processed version (upper right panel), and example of temporal traces recorded from 10 random neurons (lower panel). **(d)** Schematic showing experimental design and timeline used to record Ca^2+^ activity from D1- and D2-MSNs during sucrose or cocaine self-administration, post-abstinence seeking, and cued reinstatement. **(e)** D1-cre and D2-cre mice trained on sucrose self-administration had similar total number of NPs per session (3-way ANOVA: Time x Genotype x NP: F(11,144)= 0.909, p=0.534; Genotype x NP: F(1,144)= 0.056, p=0.814; Time x Genotype: F(11,144)= 0.888, p=0.553; Time x NP: F(11,144)= 1.973, p=0.035) and total number of sucrose pellets delivered per session (Mixed Effects 2-way ANOVA: Time x Genotype: F(11,149)= 1.724; p=0.073; Genotype: F(1,14)= 1.125, p=0.307; Time: F(4,56)= 7.201, p<0.001) over 12 days of sucrose self-administration. **(f)** D1 and D2- cre mice trained on cocaine self-administration had similar total number of NPs per session (3-way ANOVA: Time x Genotype x NP: F(11,138)= 0.776, p=0.664; Genotype x NP: F(1,138)= 1.978, p=0.162; Time x Genotype: F(11,138)= 0.701, p=0.736; Time x NP: F(11,138)= 1.540, p=0.124; and total number of cocaine infusions delivered per session (2-way ANOVA: Time x Genotype: F(11,140)= 0.700, p=0.737; Genotype: F(1,13)= 3.113, p=0.101; Time: F(11,140)= 3.938, p<0.001) over 12 days of cocaine self-administration. **(g)** Number of rewarded NPs (cued) were comparable between sucrose- and cocaine-trained D1- and D2-cre mice (2-way ANOVA: Cocaine Vs Sucrose: F(1,24)= 2.198, p=0.151, Genotype: F(1,24)= 0.051, p=0.823, and interaction: F(1,24)= 0.051, p=0.823). **(h)** Total number of NPs during first and last extinction sessions in D1- and D2-cre mice trained on either cocaine (left) or sucrose (right). **(i)** Number of rewarded NPs (cued) were higher in D1- and D2- cre mice trained on cocaine vs mice trained on sucrose (2-way ANOVA: Cocaine Vs Sucrose: F(1,23)= 5.100, p=0.0337, Genotype: F(1,23)= 0.010, p=0.923, and interaction: F(1,23)= 0.064, p=0.803). *p<0.05

### Total Ca^2+^ activity per neuron

We first quantified the total number of excitatory Ca^2+^ events for each D1- or D2-MSN. While no difference between sucrose and cocaine was found during self-administration (Figure S3a), cocaine PA seeking produced more Ca^2+^ events in D1-MSNs than sucrose, with equivalent event number between rewards in D2-MSNs (Figure S3b). For cued reinstatement the number of Ca^2+^ events in cocaine exceeded sucrose mice for both D1- and D2-MSNs (Figure S3c). Together these data indicate that for unrewarded seeking cocaine-trained mice showed more Ca^2+^ activity, especially in D1-MSNs where cocaine exceeded sucrose in both PA and reinstatement sessions (Figure S3d). In contrast, when a reward was delivered during self-administration the activity of both D1- and D2-MSNs was equivalent between cocaine and sucrose.

### Time-locked population mean of NP-induced Ca^2+^ activity

#### Rewarded seeking

We next time-locked Ca^2+^ activity in D1- or D2-MSNs to reinforced active NPs during self-administration. Heatmaps of averaged activity of D1- and D2-MSN indicated similar heterogenous response of neurons regardless of MSN subtype or reward, with some neurons activated and others inhibited around rewarded NPs (Figure 2a). Considering the heterogeneity of responses, we first averaged and compared the time-locked activity of neurons around rewarded NPs in sucrose and cocaine trained animals. Mean population activity was compared to a null distribution generated by randomly shuffling the data around the NP timestamps 1000x (95% CI was used for statistical significance). Population-averaged D1-MSN activity showed an early (0-2 sec after NP) and late (4-6 sec) excitation associated with cocaine self-administration, compared to an early excitation and late inhibition associated with sucrose self-administration (5-10 sec; Figure 2b). The comparable early excitation likely resulted from nose-poking for either reward, while the late differences between sucrose and cocaine delivery may be explained by differences in the reward delivered. Sucrose pellets were dispensed from a second port after the NP activated cue consumption, allowing us to accurately time-lock activity around NPs at the sucrose delivery port, directly associated with reward retrieval. The late decrease in D1-MSN activity corresponded to accessing sucrose pellet in the 2^nd^ port (Figure S4). Although not directly measured here, it is estimated that intravenous cocaine accesses the rodent brain within 3-7 sec^16^, supporting the possibility that the late D1-MSN excitatory response corresponded to cocaine delivery, especially considering that cocaine-induced increases in extracellular dopamine would promote depolarization of D1-MSNs^17^. Differences between sucrose and cocaine were also measured in D2-MSNs showing early excitation for cocaine and no significant change for sucrose (Figure 2c). D2-MSNs in cocaine administering mice also showed late (4-7 sec) reduction in activity, possibly a pharmacological effect of cocaine since a cocaine-induced rise in extracellular dopamine would be expected to promote hyperpolarization of D2-MSNs^17^.

**Figure 2.**
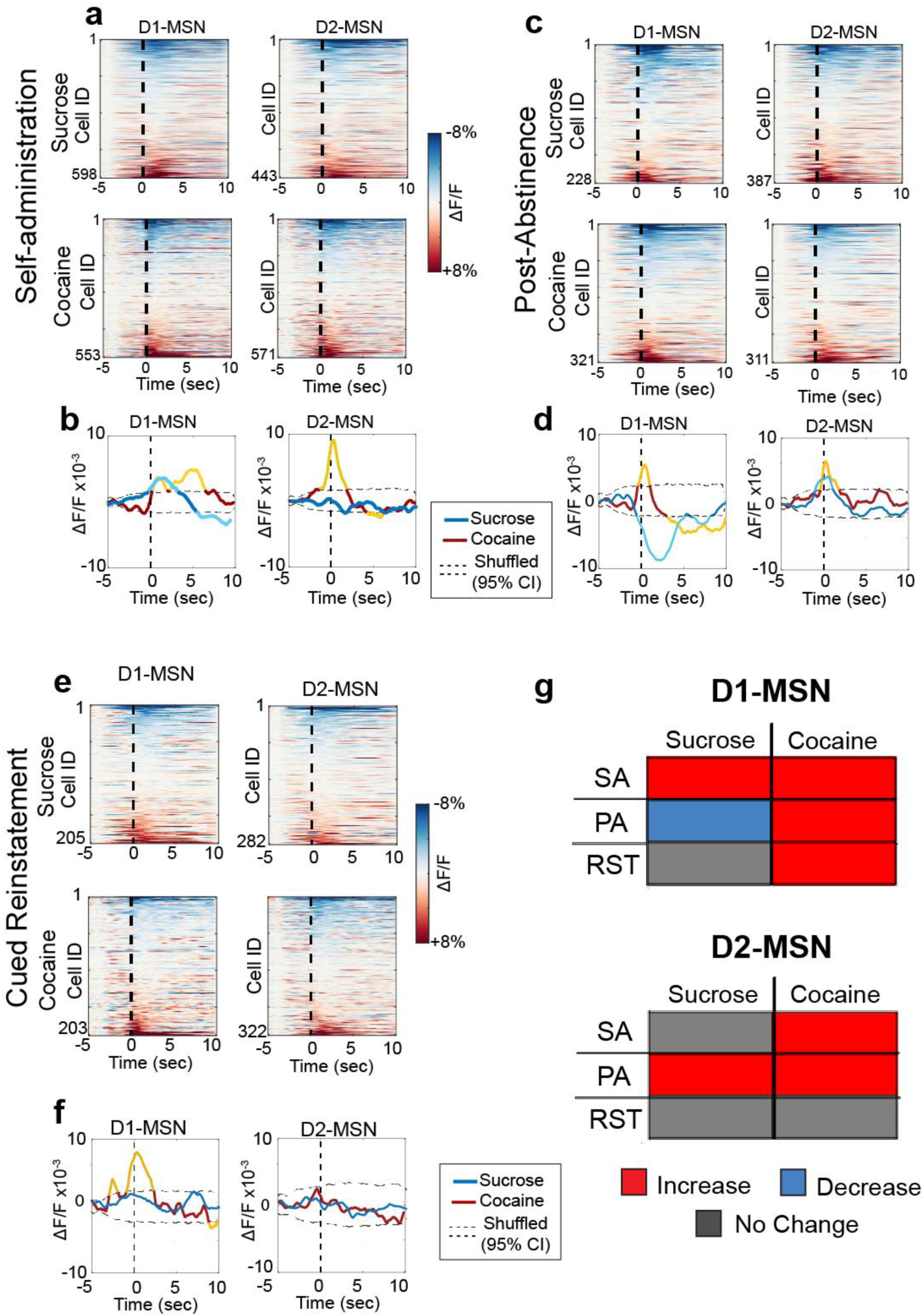
Different patterns of activity govern D1- and D2-MSN during rewarded and unrewarded cocaine/sucrose seeking. **(a)** Heatmaps representing the peristimulus histograms of the mean individual activity of recorded D1- (left) and D2-MSN (right) of mice previously trained on sucrose (upper) or cocaine (lower) around the first 10 cued active nosepoke during stable self-administration (SA7-12, 2 sessions per animal), where active nosepokes result in reward delivery and cue presentation. Dotted line represents the active NP. **(b)** Population-averaged trace activity of all recorded D1- (left) and D2- (right) MSNs during cocaine (red/yellow) and sucrose (blue/cyan) self-administration (2 sessions per animal). Dotted lines represent 95% CI of population mean activities generated by shuffling the data 1000x. Yellow and cyan lines represent population-level mean Ca^2+^ activity in cocaine or sucrose trained mice, respectively, outside the 95% CI of the shuffled distribution. **(c)** Heatmaps representing the peristimulus histograms of the mean individual activity of recorded D1- (left) and D2-MSN (right) of mice previously trained on sucrose (upper) or cocaine (lower) around the first 10 cued active nosepoke during PA seeking test, where active nosepokes resulted in cues without any associated rewards. **(d)** Population-averaged trace activity of all recorded D1- (left) and D2- (right) MSNs during cocaine (red/yellow) and sucrose (blue/cyan) PA. Yellow and cyan lines indicate mean Ca^2+^ activity outside the 95% CI. **(e)** Heatmaps representing the peristimulus histograms of the mean individual activity of recorded around the first 10 cued active NP during cued-reinstatement. **(f)** Population-averaged trace activity of all recorded D1- (left) and D2- (right) MSNs during cocaine (red/yellow) and sucrose (blue/cyan) cued-reinstatement. Yellow and cyan lines indicate statistically population-level mean Ca^2+^ activity outside the 95% CI. **(g)** Summary table showing the trends of change of population averaged activity of D1- and D2-MSN in cocaine and sucrose trained animals compared to a randomly generated shuffled distribution (red: increased activity, blue: decreased activity, Grey: no change).

#### Unrewarded seeking

D1- and D2-MSN individual activity contained heterogenous responses to NPs for unrewarded sucrose- and cocaine-seeking (Figure 2c,e). Population mean activity of all recorded neurons revealed differences between sucrose and cocaine PA (Figure 2d) and cue-reinstated seeking (Figure 2f). Population-averaged D1-MSN activity around NPs for cocaine PA rapidly increased (peak=+1 sec), while a slightly delayed decrease was observed for sucrose seeking (peak=+2 sec). Moreover, D1-MSN activity was inhibited in cocaine mice between 4-9 sec after the NP. D2-MSN activity was increased by PA seeking for both rewards peaking immediately after NP and gradually returning to baseline.

The average time-locked cue reinstated response across all neurons differed for D1-MSNs between sucrose and cocaine (Figure 2e). Cocaine cue-reinstated NPs were associated with a rapid increase in time-locked activity while no averaged time-locked activity was recorded in sucrose mice (Figure 2f). Neither sucrose nor cocaine time-locked NPs during cued reinstatement were associated with a change in overall activity in D2-MSNs (Figure 2f).

Taken together, the most striking feature in these data was that rewarded and unrewarded cocaine seeking activity was associated with consistent excitatory responses in D1-MSNs (Figure 2g). In contrast, D1-MSN responses to sucrose seeking activity varied depending on the session and reward availability, from increase to decrease to no change.

### Subpopulations of MSNs time-locked to NPs

#### Rewarded Seeking

Due to the heterogeneity of the neuronal responses recorded around seeking NPs, we clustered individual neurons into subpopulations based on their time-locked peri-NP activity over the 5 sec before and 10 sec after rewarded NPs. Individual neurons were parsed into excited, inhibited or not time-locked, by comparing their mean activity across all trials to a null distribution generated by shuffling the Ca^2+^ activity around behavioral responses (1000x iterations, 95% confidence interval; Figure 3a,b and Figures S5-6; see Methods for details). The subpopulations of neurons were compared between cocaine- and sucrose-trained mice across the behavioral protocol in Figure 1d, including during self-administration, PA seeking, and cued reinstatement.

**Figure 3.**
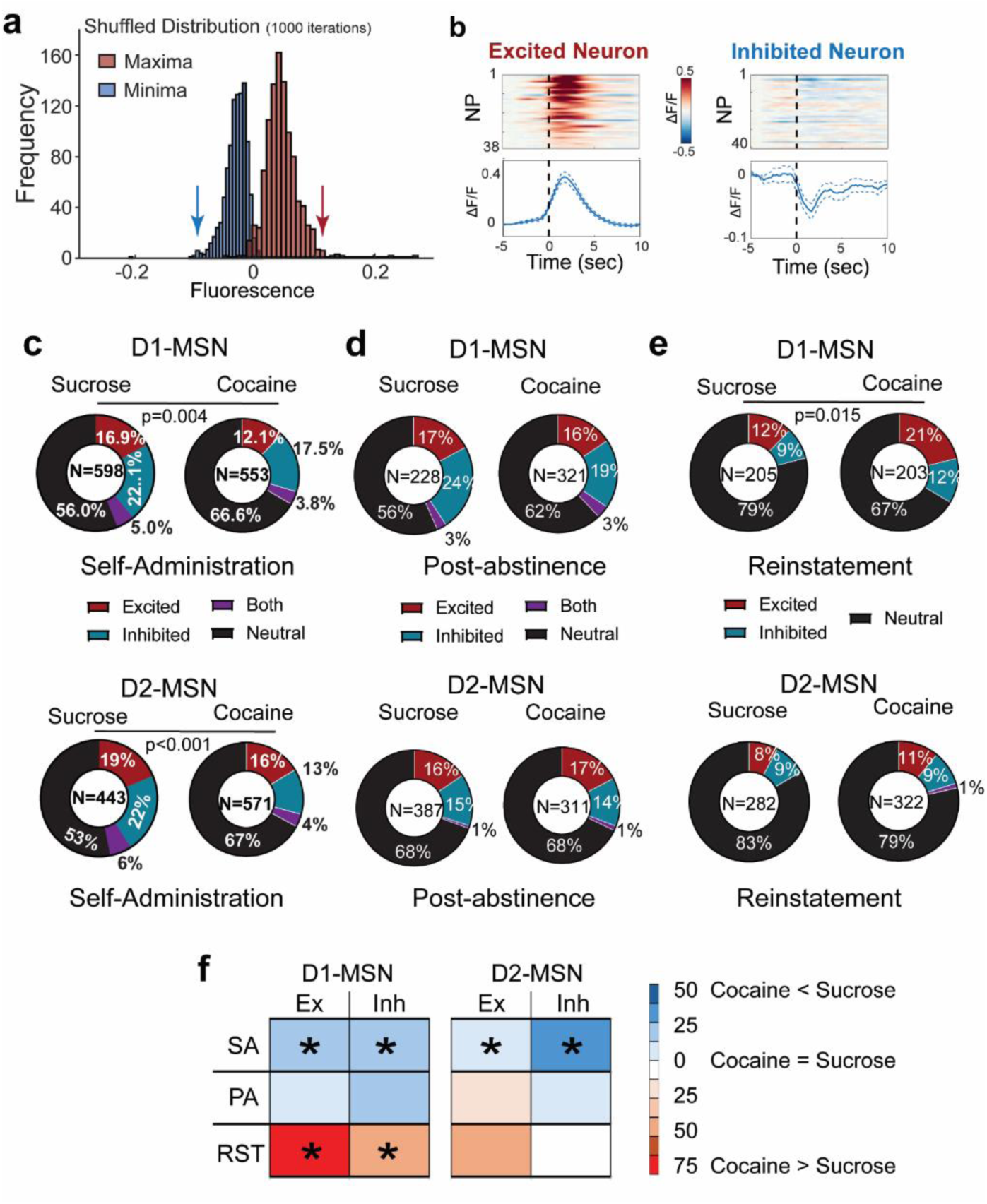
Subpopulations of D1- and D2-MSN time-locked to reward seeking activity. **(a)** An example of the distribution of the maxima and minima of mean neuronal activity around nosepoke (5 seconds before to 10 seconds after) generated after shuffling the calcium trace of each neuron around behavioral data points (1000x). If the maximum or minimum activity of the real data was higher or lower than 97.5% of the maximum (right) or minimum (left), respectively, the neuron was considered to be time-locked excited or inhibited. Arrows indicate the 97.5% maximum and minimum thresholds. **(b)** Examples of peri-event activity histograms of time-locked excited (left) and time-locked inhibited (right) neurons. Each row of the heatmaps represents an individual trial, i.e. a cued nosepoke. Activity traces in the lower panel represent mean ± SEM activity across all trials. **(c)** Pie charts representing proportions of excited (red) and inhibited (blue) neurons during cocaine or sucrose self-administration show higher proportions of time-locked excited and inhibited neurons in sucrose vs cocaine trained mice. **(d)** Pie charts representing proportions of excited (red) and inhibited (blue) neurons during cocaine or sucrose post-abstinence seeking tests show no difference between both groups. **(e)** Pie charts representing proportions of excited (red) and inhibited (blue) neurons during cocaine or sucrose cued reinstatement seeking tests show no difference between both groups. Cocaine and sucrose were compared using a Chi square test. **(f)** Summary table showing differences in distribution of excited/inhibited timelocked neurons between during different phases of cocaine and sucrose seeking. Coding = %cocaine neurons/ %sucrose neurons within each subpopulation. *p< 0.05 based on Chi square test comparing sucrose and cocaine subpopulations within each behavioral trial

During rewarded self-administration, sucrose NPs yielded a higher proportion of time-locked excited and inhibited subpopulations of D1- and D2-MSNs than cocaine NPs (Figure 3c). Notably, the activity of excited D1-MSNs during cocaine self-administration showed a bimodal distribution (figure S5), akin to that of the mean population activity (Figure 2c). The bimodal distribution is consistent with two different populations of excited D1-MSNs, the first activated between 0- 2seconds, and the second activated between 3-7 seconds, which supports the earlier conclusion that the two peaks may be due to an early cue response, followed by a delayed pharmacological cocaine-mediated excitatory response.

#### Unrewarded Seeking

During PA seeking, the proportion of subpopulations of D1- and D2-MSNs showing excitation, inhibition, both or not time-locked were comparable in size between sucrose- and cocaine-trained mice (Figure 3d). Notably, cue reinstated cocaine seeking recruited more overall time-locked D1-MSNs than sucrose seeking, which was explained predominantly by a nearly 2-fold increase in excited D1-MSNs (Figure 3e,f). In general, time-locked excited D1- MSNs were more excited and more synchronized around NPs during unrewarded cocaine seeking sessions compared to sucrose (Figure S5), while time-locked inhibited neurons were more inhibited during sucrose seeking sessions compared to cocaine (Figure S6). On the other hand, the proportions of D2-MSN showed no differences between sucrose or cocaine during unrewarded post-abstinence or reinstated seeking during (Figure 3d,e).

Taken together (Figure 3f), these data indicate that during self-administration, when the reward is available, sucrose reinforcement is recruiting more excited and inhibited D1- and D2-MSNs, perhaps contributing to the fact that when given a choice, rodents prefer food over cocaine reward^4,15^ (however, see^18^). When the reward is omitted during cued reinstatement test, cocaine associated NPs and cues recruit more excited and inhibited D1-MSNs than sucrose associated NP/cues.

### Stability of time-locked activity within a seeking session

We tracked time-locked individual neurons across the course of a seeking session to determine if the responses of neuronal subpopulations were stable or different neurons were recruited to the time-locked excited and inhibited subpopulations of MSNs. Stability was estimated by time-locking to odd- or even-numbered NPs, and if an MSN was time-locked to both sets of events, it was considered stable (Figure 4a shows examples of stable and unstable neurons).

**Figure 4.**
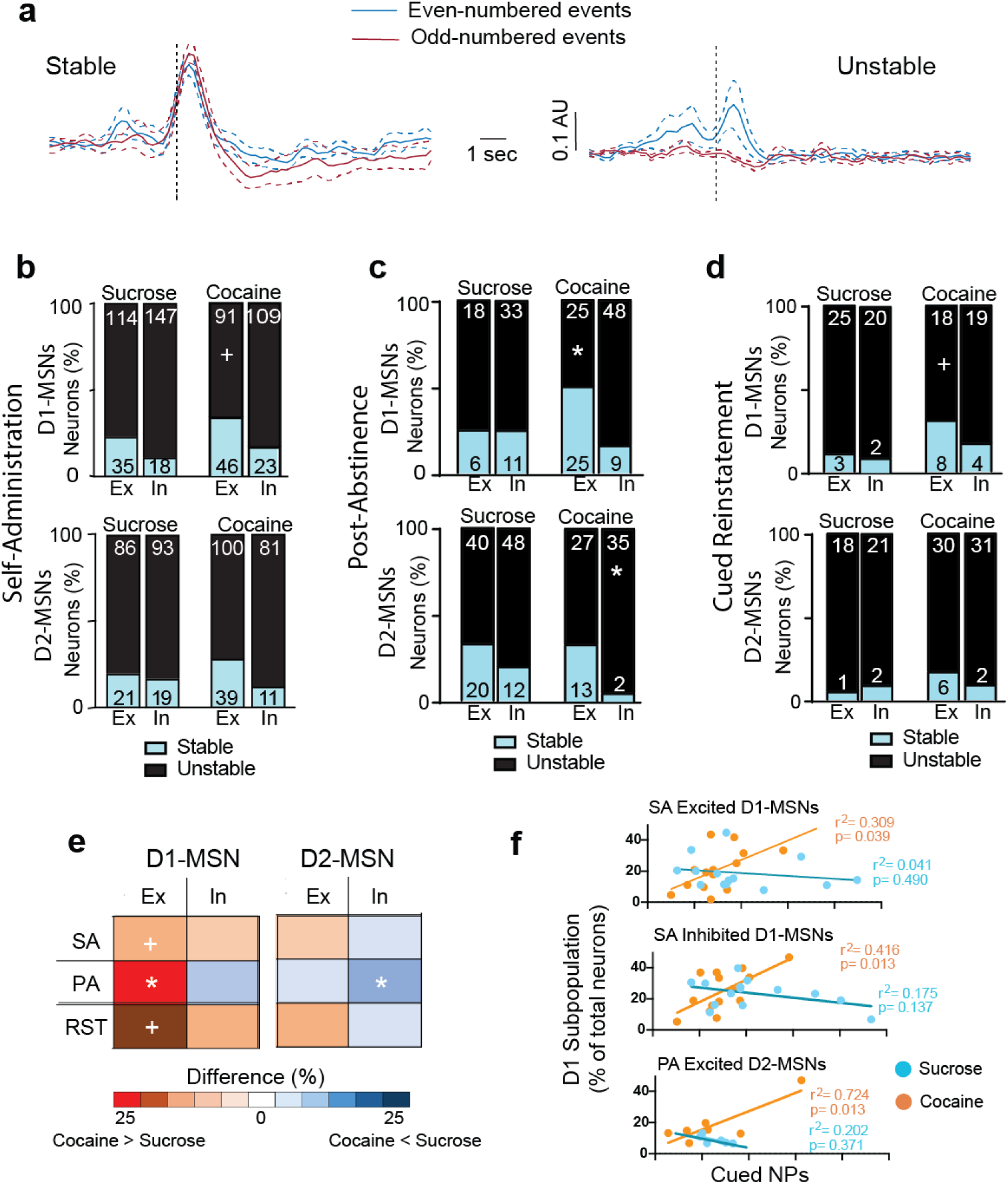
Cocaine seeking is characterized by stable within session excited D1-MSN population. **(a)** Example traces of stable and unstable neurons showing respectively consistent and inconsistent mean increase in activity during odd- and even-numbered events. (Red: mean activity around odd-numbered events; blue: mean activity around even numbered events; dotted lines represent the standard error). **(b)** Bar graphs comparing the stability (blue – stable, black – unstable) of excited (Ex) and inhibited (In) D1-MSN (left) and D2-MSN (right) ensembles during stable self-administration between sucrose- and cocaine-trained animals by comparing time-locking to odd-numbered events to even-numbered events. (* p<0.05 cocaine vs sucrose; + p<0.10; see Table S1 for all Chi^2^ values). **(c)** Same graph as (b) for post-abstinence test. **(d)** Same graph as (b) for cued-reinstatement. **(e)** Summary table showing percentage of stable neurons within each ensemble during cocaine/sucrose self-administration (SA), post-abstinence seeking (PA), or cued reinstatement (RST). (* p<0.05 comparing ensembles between cocaine sessions and sucrose sessions; + p<0.10; see Table S1 for Chi^2^ values). **(f)** Excited D1-MSN significantly predict total number of rewarded and PA NPs in cocaine-trained mice (see Table S2 for r^2^ values).

#### Stability of time-locked activity during reward seeking

For either self-administered rewards, the proportion of stable excited D1-MSNs trended (p= 0.059) higher in cocaine than sucrose self-administration. No other differences in stability were evident between sucrose and cocaine across the subpopulations.

#### Stability of time-locked activity during unrewarded seeking

We performed the same subpopulation stability analyses for both the PA seeking and cue reinstatement. During the PA session the odd/even analysis for stability showed two major differences (Figure 4c). Excited D1-MSNs in cocaine mice were more stable than sucrose excited D1-MSNs. Also, inhibited D2-MSNs in cocaine mice were remarkably unstable and showed less stability than sucrose inhibited D2-MSNs.

The odd/even analysis for cued reinstatement showed that excited D1-MSNs trended (p= 0.068) towards being more stable in cocaine compared to sucrose mice (Figure 4d). No other differences in stability between sucrose and cocaine were evident between sucrose and cocaine across the subpopulations.

The patterns of subgroup stability across the three different seeking sessions are summarized in Figure 4e, which illustrates the percent difference in stable MSNs between cocaine and sucrose. The primary result is that excited D1-MSNs are more stable in response to active seeking in cocaine-compared with sucrose-trained mice, regardless of the type of seeking session (self-administration, PA or cued reinstatement). Also, relative to sucrose, cocaine mice had fewer stable inhibited D2-MSNs in the PA session.

#### Correlations between time-locked subpopulations and NPs

We examined if the neuronal subpopulations were statistically associated with cued active NPs by linear regression comparisons between cued NPs and the percent of neurons in each subgroup. Combining D1- and D2-MSNs revealed that cocaine excited MSNs were correlated with NPs during self-administration and PA seeking and cocaine inhibited MSNs correlated with NPs during self-administration (Table S2). When the MSNs were separated into the excited D1- and D2-MSN subpopulations, only cocaine excited D1-MSNs remained correlated with NPs during self-administration and PA (Figure 4f, Table S2). Similarly, only inhibited D1-MSNs were correlated with cocaine NPs (Figure 4f). No correlations were observed during cued-reinstatement, nor in sucrose seeking for any subpopulation of MSNs (Figure 4f; Table S2).

### Decoding NP initiation using MSN subpopulations

Neuronal responses preceding NPs may represent a motivational signal to seek the reward. We used machine learning (Figure S7) to determine if Ca^2+^ data from time-locked subpopulations of D1- and D2-MSNs could decode the initiation of an NP (activity during the 5 sec preceding NP). The decoding accuracy in shuffled datasets of random events were used as control and were subtracted from the quantified time-locked data in each subgroup for each animal in the study.

Rewarded NPs during self-administration were decoded by excited D1-MSNs in sucrose and cocaine mice and excited D2-MSNs only in cocaine mice (Figure 5). Only in cocaine mice did inhibited D1-MSNS effectively decode NPs. Surprisingly, MSNs that were not time-locked decoded sucrose, but not cocaine rewarded NPs for both D1- and D2-MSNs. Akin to rewarded NPs, for both sucrose and cocaine PA seeking excited D1-MSNs predicted NPs and inhibited D1-MSNs decoded only cocaine NPs (Figure 5a). Excited D2-MSNs also predicted PA NPs for cocaine, but not sucrose (Figure 5b). During cued reinstatement excited D1-MSNs decoded NPs only in cocaine mice and no other subgroup of D1- or D2-MSNs successfully decoded sucrose or cocaine NPs. Figure 5c summarizes the capacity of subpopulations of MSNs to decode cocaine and sucrose NPs in all three behavioral sessions. Similar to the above observations of subpopulation stability and overall activity (Figures 3f, 4e), only excited D1-MSNs decoded NPs in all three cocaine seeking sessions. Also, it was surprising that for sucrose rewarded NPs both the D1- and D2-MSNs that were not time-locked predicted the NPs. This argues that other subpopulations not identified using our time-locking algorithm may be contributing to sucrose but not cocaine self-administration.

**Figure 5.**
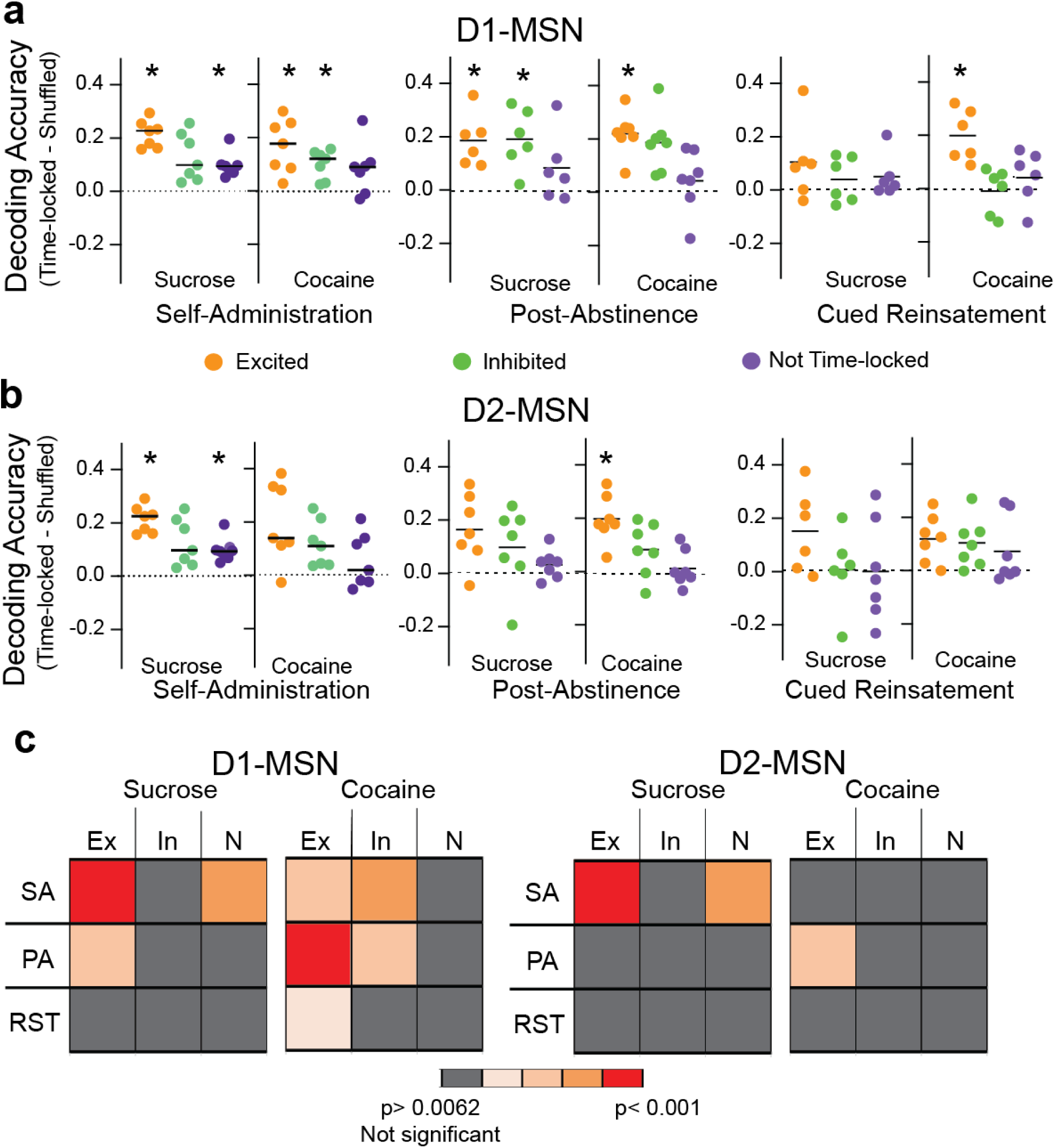
Decoding nose poking by different MSN subpopulations. **(a)** Nose poke decoding accuracy using the neuronal data of the three types of D1-MSN activity patterns during self-administration (left), post-abstinence (middle) and cued-reinstatement (right). * p< 0.0062, using a Student’s t-test comparing measured data to shuffled data points with the p values corrected for multiple comparisons using a false discovery rate, q= 0.02. Table S3 contains all t-test and probability values. **(b)** Nose poke decoding accuracy using the neuronal data of the three types of D2-MSN activity patterns. * p< 0.0062, using a Students t-test comparing measured data to shuffled data points with the p values corrected for multiple comparisons using a false discovery rate of q= 0.02. **(c)** Summary table showing the relative NP decoding accuracy of subtypes of D1- and D2-MSN between sucrose and cocaine trained mice. Ex- excited, In-inhibited, N- not time-locked

### Stability of time-locked activity between self-administration sessions

Lastly, we determined if time-locked subpopulations of MSNs were stable longitudinally between different rewarded self-administration sessions, and if the same neurons associated with cocaine self-administration had similar responding in a subsequent unrewarded PA session. Neurons between multiple sessions were tracked using a previously established nearest-neighbor cell registration method^19^ (Figure 6a). The same D1-MSN and D2-MSN were tracked between two late sucrose and cocaine self-administration sessions (Figure 6b,c) where mouse self-administration was stable and reliable (day 7-9 versus 10-12, respectively). Neurons were termed stable if they showed similar time-locked responses during the first and second self-administration sessions. There was no difference in stability between sucrose and cocaine rewarded seeking in either of the D1- or D2-MSN subpopulations (Figure 6b,c; Table S4 for statistical values). To verify if neuronal responses are preserved across multiple self-administration sessions, we used machine learning to train an SVM model on tracked neurons from an early self-administration session (day 7-9) to decode NPs from a later self-administration session (days 10-12). Tracked D1-MSNs, but not D2-MSNs, from one self-administration session successfully decoded seeking behavior from the other session in cocaine and sucrose trained animals (Figure 6d; Table S5 for statistical values). This indicates that a relatively stable neuronal representation of D1-MSNs was formed during self-administration in both reward subtypes.

**Figure 6.**
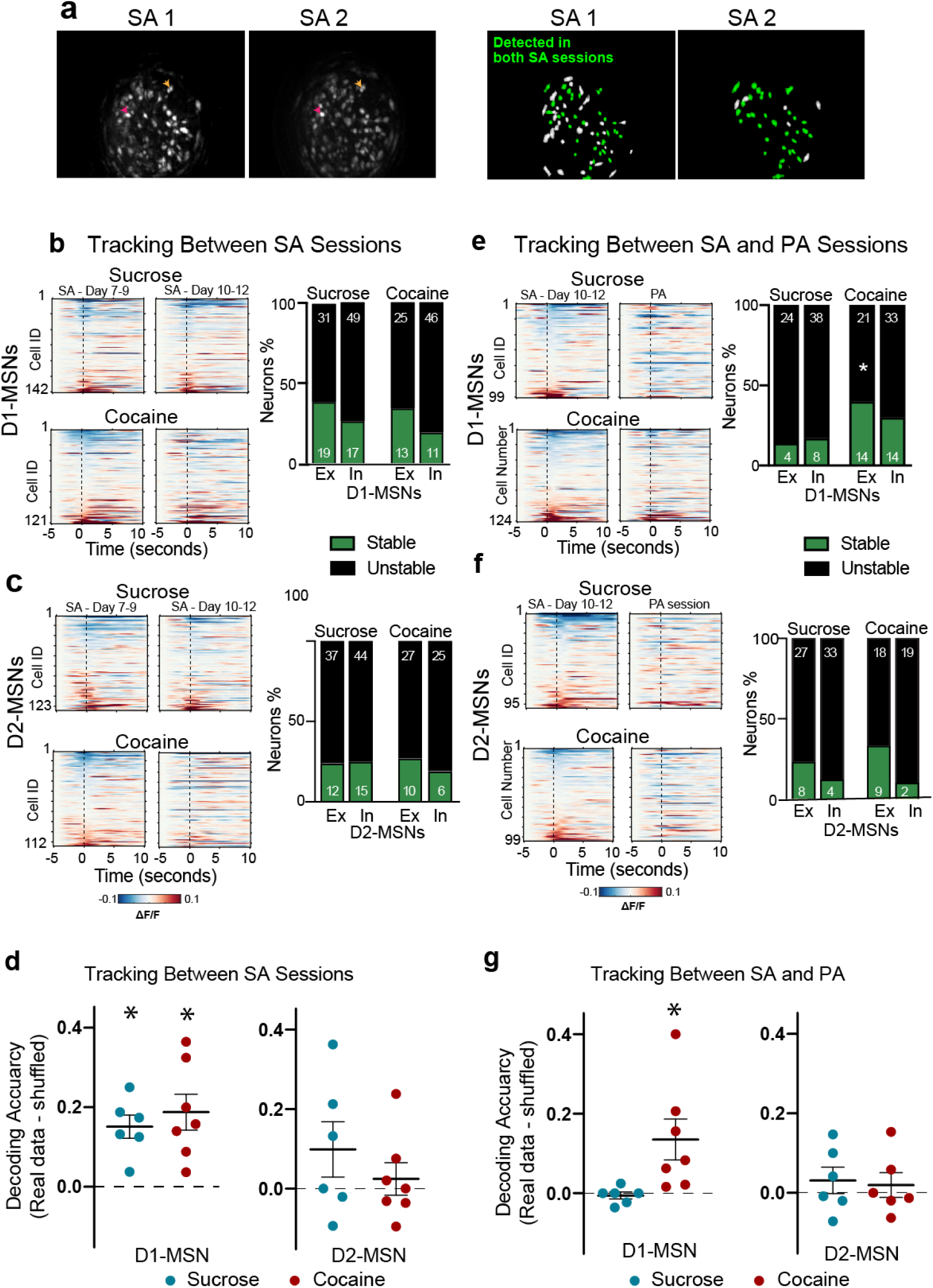
Tracking the same neurons across self-administration (SA) to PA sessions shows a higher stability of responding in the excited D1-MSN subpopulation. **(a)** Left: Maximal intensity projections of recordings from two stable self-administration sessions recorded within the same animal, showing similar fields of view; Right: schematic of the spatial footprints of all identified neurons (green+white) in each session. The neurons that were successfully tracked over both sessions are colored green. The neurons that were only visualized in one session but not the other are colored white. Colored arrows point towards to examples of neurons detected in both sessions. **(b)** Left: Heatmaps representing the mean activity of D1-MSNs longitudinally tracked between two stable cocaine or sucrose self-administration sessions. Each row represents one neuron tracked across both sessions. Right: Bar graphs comparing the stability (green– stable, black– unstable) of excitatory and inhibitory D1-MSNs across two self-administration sessions. **(c)** Heatmaps representing the mean activity of D2-MSNs longitudinally tracked and stability of neuronal activity between two cocaine or sucrose self-administration sessions (SA 8/9 vs SA 10/11). There were no differences in proportion of stable neuron subpopulations between sucrose and cocaine. **(d)** Decoding of the second SA session by training on the first SA session. *p<0.05, paired t-test comparing each subpopulation with its shuffled distribution, Table S5 for all t and p values. **(e)** Heatmaps representing the mean activity of D1-MSNs longitudinally tracked and stability of neuronal activity between two cocaine or sucrose SA and PA sessions. Only stable excited D1-MSNs differed between cocaine and saline (Chi^2^ and p values in Table S4). **(f)** Heatmaps representing the mean activity of D2-MSNs longitudinally tracked and stability of neuronal activity between cocaine (lower) or sucrose (upper) self-administration (SA 10/11) and PA sessions. No differences between sucrose and cocaine were found (Chi^2^ and p values in Table S4). **(g)** Scatter plot showing the decoding accuracy of SVM model trained on the neuronal activity of D1- or D2-MSNs during the SA session and subsequently used to decode NPs during the PA session. Only cocaine D1-MSNs decoded PA NPs (t and p values in Table S5).

### Stability of time-locked activity between rewarded and unrewarded sessions

One key aspect of drug seeking behavior is the long-lasting associations made between drug seeking and drug-associated cues, whereby associated cues gain salience to drive behavior despite an extended period of abstinence. We asked if the neuronal representations that we recorded during cocaine and sucrose self-administration were carried forward into unrewarded PA seeking sessions, with the overarching hypothesis that stable neuronal representations are likely responsible for the prepotent seeking for cocaine over sucrose. We longitudinally tracked neurons between the last self-administration session (day 10-12) into the PA session conducted after 7-10 days of abstinence. Akin to the within session stability measurements (Figure 4), excited cocaine D1-MSNs were more stable than sucrose D1-MSNs across the two sessions (Figure 6e; Table S4). We also trained machine learning models on the recorded data from tracked neurons from self-administration session to decode NPs during PA sessions. Interestingly, only cocaine D1-MSNs that tracked from self-administration to PA sessions accurately decoded seeking in the PA session (Figure 6g; Table S5 for statistical values), indicating a stable and persistent representation of cocaine seeking compared to sucrose seeking despite an extended period of abstinence.

Together with the within session stability data (Figure 4) and the overall responsiveness time-locked D1- and D2-MSNs (Figure 2), the across-session stability further emphasizes the role of greater excited D1-MSN stability in cocaine versus sucrose trained mice across the rewarded and unrewarded (PA) seeking modalities, indicative of a stable long-term ensemble consistently recruited during cocaine seeking events.

## DISCUSSION

We used a variety of analytical approaches to determine if cocaine and sucrose seeking involved neuronal activity in distinct subpopulations NAcore MSNs, with the overarching hypothesis that any differences discovered might contribute to why individuals with cocaine use disorder are more motivated by cues predicting cocaine over natural rewards. Using single cell Ca^2+^ imaging as a measure of neuronal activity allowed us to quantify activity in specific subpopulations of D1- and D2- MSNs and to longitudinally track the subpopulations within and between rewarded and unrewarded seeking^19^. The most striking and consistent distinction we discovered between sucrose and cocaine seeking was that excited D1-MSNs show greater overall activity, stability and capacity to decode cocaine seeking behavior compared to sucrose seeking. This difference in D1-MSN activity between cocaine and sucrose seeking was most evident during unrewarded cocaine seeking (PA and cue reinstated seeking). In contrast, similar patterns of activity in D1- and D2- MSNs were often manifested by sucrose and cocaine seeking when the reward was available during self-administration.

During rewarded self-administration sessions, both reward types were associated with comparable peri-NP excitation of D1-MSNs, similar heterogeneity of responses across excited/inhibited subpopulations, and comparable decoding accuracy of D1- and D2- neuronal data to predict NPs. However, a difference between sucrose and cocaine in D1-MSN activity was observed between 5- 10 sec after the rewarded NP that likely corresponded to receipt of the reward. Cocaine showed an delayed excitatory response associated with intravenous cocaine delivery that contrasted with a delayed inhibitory response associated with sucrose reward consumption. Interestingly, distinct and non-overlapping subpopulations of excited D1-MSNs were associated with the immediate NP and the subsequent delayed response to reward, suggesting different neuronal ensembles and possibly brain circuits involved in each response^11^. While the cocaine-associated delayed increase is consistent with cocaine’s increase in dopamine stimulation of D1-dopamine receptors^20^, the findings that sucrose retrieval was associated with a decrease rather than an increase in D1-MSNs activity of the NAcore was surprising but consistent with previous in vivo electrophysiological recordings from the nucleus accumbens during food consumption^21,22^.

While excited D1-MSNs were also associated with rewarded sucrose seeking by some measures, the activity was not as consistent as with cocaine for unrewarded seeking. For example, excited D1-MSN activity during unrewarded cocaine sessions was notably more robust, of higher amplitude and more synchronized around seeking activity (Figure S5). Also, our different measures of neuronal stability showed a higher consistency within the excited D1-MSN subpopulation in cocaine-trained animals compared to sucrose in unrewarded sessions. Furthermore, excited D1-MSNs decoded and predicted NPs only in cocaine, not sucrose cue seeking. Together, these findings indicate that an excited D1-MSN subpopulation formed a consistent and stable ensemble mobilized by unrewarded cocaine seeking. Importantly, while this ensemble was formed and stable for both sucrose and cocaine self-administration, only the cocaine excited D1-MSN ensemble persisted into unrewarded PA seeking. In contrast, the parallel ensemble of excited D1-MSNs present during sucrose self-administration did not reform during PA seeking for sucrose. In conclusion, an excited D1-MSN ensemble formed during self-administration of either cocaine or sucrose, but propagated to unrewarded seeking only in cocaine-trained mice. This conclusion is buttressed by finding that only excited D1-MSN activity correlated with the intensity of cocaine PA seeking, while no subpopulation of MSNs correlated with sucrose behavioral seeking. Moreover, we replicated our previous findings and the findings of others^4,15^, that cue reinstated seeking is greater for cocaine than sucrose cues.

The partial similarity between the excited D1-MSN subpopulations during rewarded cocaine or sucrose seeking and the striking difference in this subpopulation during unrewarded sucrose versus cocaine seeking, raises an important question. What mediates the propagation of the excited D1- MSN subpopulation from rewarded into unrewarded seeking only in cocaine mice? Indicative of potentiated D1-MSNs synapses, non-contingent or self-administered cocaine infusions produce enduring increases in dendritic spine density and spine head diameter selectively in accumbens D1-MSNs^23–25^. This is paralleled by an increase in AMPA glutamate receptor mediated currents after extended withdrawal from cocaine self-administration^26–28^. Importantly, enduring potentiation is found selectively in D1-MSNs after abstinence from cocaine, not sucrose self-administration^29^. While the enduring adaptations induced by cocaine, not sucrose, demonstrate a mechanism whereby D1-MSN excitatory responses to glutamate release are potentiated for weeks after discontinuing drug use, measurements of synaptic potentiation during unrewarded seeking reveal that cocaine, but not sucrose cues elicit further potentiation that is transiently expressed only for the duration of a seeking session^30^. For example, ex vivo measures of synaptic potentiation, including increased AMPA/NMDA ratio and dendritic spine head diameter, are transiently elevated only in D1-MSNs during cued cocaine, not sucrose seeking^29,31^. This is paralleled by increased extracellular matrix signaling through b3-integrins selectively in D1-MSNs^32–34^ and decreased astroglial synaptic proximity during cued seeking for cocaine or heroin, not sucrose^35^. Combined these data are highly consistent with our Ca^2+^ activity measurements revealing an ensemble of excited D1-MSNs associated with NPs for cocaine PA and cued reinstatement, but not for unrewarded sucrose seeking. Furthermore, the data offer a mechanism by which this occurs, via both enduring synaptic potentiation in withdrawal and transient synaptic potentiation that depends on temporary adaptations in astroglial morphology and signaling in the extracellular matrix.

A role for D1-MSNs as key regulators of reward seeking is consistent with a variety of optogenetic^36^ and DREADD stimulation and inhibition^31,37,38^ studies concluding that activity in D1-MSNs is sufficient and necessary for rewarded and unrewarded cue seeking^4,30,31^. Conversely, these studies generally conclude that activity in D2-MSNs serves an opposing function to reduce reward seeking^30,31,37,39^. However, optogenetic and DREADD studies indiscriminately stimulate or inhibit D1- or D2-MSNs, which does not reflect the electrophysiological and imaging observations that subpopulations of MSNs can be active and inhibited during a seeking event regardless of genotype^5,7^. In accord with the importance of this caveat, the functional dichotomy between D1- and D2-MSNs is being challenged by studies showing a more nuanced collaborative role between MSN subpopulations^40–42^. For example, if animals undergoing operant food seeking are tested during periods of reward unavailability, fiber photometry recordings reveal increased Ca^2+^ activity in both MSN subtypes, and inhibiting either subtype results in associated increases of unproductive reward seeking^43^. Also, 2-photon imaging from the shell subcompartment of nucleus accumbens shows that distinct functional clusters of both D1- and D2-MSNs encode the intensity of sucrose rewards versus consummatory (i.e. licking) behavior^44^. Furthermore, a recent examination of how valence of a conditioned stimulus controls operant responding concluded that D1- and D2-MSNs collaborate to provide information regarding specific valence-independent associative learning^45^. Finally, the dichotomous model of D1- and D2-MSN functioning fails to account for further genetic heterogeneity within each subtype, such as a recently identified subpopulation of D1-MSNs expressing tachykinin-2 that negatively regulates cocaine seeking behavior^46^. While our study reveals both excited and inhibited subpopulations of D1- and D2-MSNs in the NAcore can be associated with both cocaine and sucrose seeking, only the excited D1-MSN population was stable and predictive of unrewarded cocaine, not sucrose seeking. Nonetheless, these data do not discount a role for other subpopulations of MSNs contributing to both cocaine and sucrose seeking. Consistent with this idea, there is ∼25% overlap between neurons labeled for the IEG c-fos during cue-induced cocaine and sucrose seeking^4^.

Our experiments show that cocaine seeking is associated with a stable ensemble of D1-MSNs activity within rewarded seeking sessions that persists for at least 10 days of abstinence into a post-abstinence seeking session. Stability of individual neuronal activity within the 120 min rewarded and unrewarded seeking sessions as well as longitudinally after 10 days of abstinence between a rewarded and unrewarded sessions strongly argues that a discrete ensemble of excited D1-MSNs is formed during self-administration and becomes critical for unrewarded cocaine seeking. Conversely, the relative instability and poor decoding of time-locked MSN responses during unrewarded sucrose seeking is consistent with network level neuronal encoding that likely involves recruitment of shifting populations of D1- and D2-MSNs to guide behavior. This interpretation resonates with behavioral studies directly comparing cocaine and sucrose seeking that generally show neuronal activity in the NAcore is obligatory for cued reinstatement of cocaine, but not sucrose seeking^31,47,48^. Thus, sucrose seeking involves a more distributed network in the brain that can function in the absence of NAcore neurons, possibly reflecting the survival necessity of food seeking^10^. In contrast, cocaine seeking is associated with drug-induced enduring and transient neuroadaptations selective for NAcore D1-MSNs that favor the formation of an excited D1-MSN ensemble underpinning cocaine NPs^29,30,49^. Based on our observations of a cocaine, not sucrose seeking ensemble of excited D1-MSNs and other findings identifying long-lasting and cue-induced transient adaptations by cocaine over sucrose seeking, we hypothesize it is the formation of this ensemble that causes cocaine cues to more potently drive seeking compared to sucrose cues^4,50,51^, and contributes to why individuals suffering cocaine use disorder seek cocaine in response to cocaine-associated stimuli in preference to seeking natural rewards.

## ACKNOWLEDGMENTS

This work was supported by the National Institute of Health (DA051159 to R.M.C., and DA046373 and DA012513 to P.W.K.).

## Author Contributions

Conceptualization, R.M.C. and P.W.K.; Methodology, R.M.C. and P.W.K.; Investigation, R.M.C., A.T., J.H., and C.C.; Software and Data Curation: R.M.C. Writing – Original Draft, R.M.C. and P.W.K.; Writing – Review & Editing, R.M.C. and P.W.K.; Funding Acquisition, R.M.C. and P.W.K.; Visualization: R.M.C. and P.W.K.

## SUPPLEMENTAL INFORMATION

Document S1. Supplemental figures (S1-S6)

Document S2. Supplemental tables (T1-T5)

## Methods

### Resource Availability

Lead Contact

Further information and requests for resources and reagents should be directed and will be fulfilled by the lead contact, Reda M. Chalhoub

#### Materials Availability

This study did not generate new unique reagents

#### Data and code availability

Data reported in this paper, as well as custom MATLAB codes used for the analysis, are available from the lead contact upon reasonable request.

Any additional information required to reanalyze the data reported in this paper is available from the lead contact upon request.

### Experimental model and subject details

#### Animals

Adult male and female transgenic D1-cre (129S6.FVB(B6)-Tg(Drd1a-cre)AGsc/KndlJ, Jax Strain #:028298), D2-cre (B6.129S4(FVB)-Drd2tm1.1Mrub/J, Jax Strain #:010687), and A2A-cre (B6;129-Adora2atm1Dyj/J, Jax Strain #:010687) mice were used in the experiments. All mice were bred in-house and periodically outcrossed with wild-type C57/BL/6J mice to maintain genetic diversity. Mice were group housed before surgeries, and single-housed after surgeries to avoid interference with behavioral experiments and minimize the risk of infection or injury resulting from the interaction with cage mates during the recovery period. After surgical procedures were performed, animals were kept in a reverse light cycle (12-hour dark/12-hour light) under controlled temperature and humidity settings. All behavioral experiments were run in the Dark Phase. Mice were at least 12 weeks old and weigh 20g prior to their first surgery. All experiments and procedures were performed in compliance with the guidelines of the institutional Animal Care and Use Committee at the Medical University of South Carolina.

### Method details

#### Surgeries

All surgical procedures were conducted under isoflurane anesthesia (induction: 5%, maintenance: 1-2%, flow rate: 0.2 L/min).

##### Surgeries for endoscopic calcium imaging

After anesthesia induction and preparation and sterilization of the scalp, a small incision was made to expose the skull surface. A dental drill (5mm diameter) was used to create one craniotomy hole over the left nucleus accumbens (AP: +1.6 mm, ML: +1.1 mm relative to Bregma). To achieve cell-type specific calcium imaging, mice were stereotaxically injected with an adeno-associated viral vector encoding a double-floxed inverted orientation GCaMP6f (pAAV-Syn-Flex-GCaMP6f-WPRE-SV40, titer: ∼ 1 × 10^13^ GC/mL, acquired from Addgene (#100833)) using Nanoject III (Drummond Scientific, total volume = 750 nL, rate of injection: 50 nL/min, total duration 10 mins). Four weeks following viral injection, a lens with an integrated baseplate (Inscopix, 0.6 mm diameter, 7.3mm length) was slowly lowered (0.2mm/min) over the same location, under active miniscope visualization and guidance, until a clear and bright plane was visualized, indicating proper expression of the virus and alignment of the lens. The chosen z-plane was constrained to be between −4 and −5 to ensure placement in the nucleus accumbens core. The lens and the baseplate were secured in place using two stainless steel screws (to the skull surface) and dental cement (1:5 mix of black and transparent acrylic cement). The animal’s lens was covered with a magnetic cap (Inscopix) to protect it from scratches and physical damage. All mice were weekly monitored through imaging trials over 4 weeks after the surgery to detect single-cell neuronal activity; if no single neurons were visualized, mice were euthanized and excluded from the study.

##### Catheter Surgeries

For cocaine self-administration in mice, mice were implanted with a custom-made indwelling jugular catheter connected to a back-mounted guide cannula, as previously described^52,53^. In brief, a 12-mm catheter tip was subcutaneously passed from the back and inserted in the right jugular vein through a 1cm mid-scapular incision. The catheter tip was sutured in place to the vein and subcutaneous adipose tissue, while the entry port cannula was secured to the back. All mice received a perioperative analgesic injection of Carprofen (5mg/kg, subcutaneously) and Cefazolin (200 mg/kg, iv) to avoid infections, and were then treated daily with carprofen (5 mg/kg, subcutaneously) and topical antibacterial ointment for 3 days post-operatively. Incision sites were treated with topical triple antibiotic ointment and Catheters were flushed daily with heparin (100 units/mL) to prevent catheter occlusion and maintain patency. Patency of the catheters was assessed by injecting the mice with 0.01-0.02mL of Brevital (2mg/mL): an immediate loss of muscle tone within 0-5 seconds following the injection indicated a positive test and a patent catheter. Mice with non-patent catheters at any point during self-administration were excluded, and catheter patency was no longer assessed after the start of home-cage forced abstinence.

#### Self-administration

After recovering from their respective surgical procedures, mice were switched to a restricted food diet (∼3g of chow daily) to maintain their weight at 90% of their original weight. Self-administration was performed in an operative box (Med Associates, Inc) that contains: a house light, two nosepoke ports, an availability light over the active nosepoke, a syringe pump (used for cocaine self-administration), and a sucrose pellet dispenser connected to a tall food pellet receptacle (used during sucrose self-administration).

##### Sucrose Self-Administration

Each session started with a 15-minute “baseline” period during which the house light was off and the nosepokes did not result in sucrose delivery, followed by 120-minutes of sucrose self-administration. The “baseline” period was used to adapt the animal to the headcap connections (optical fiber cables for experiments involving optogenetics or miniscope for experiments involving calcium imaging). A maximum of 100 sucrose pellets per session was allowed to prevent an overdose. After self-administration, mice undergo abstinence from sucrose self-administration for 10 days in their home cage during which they were handled daily. After the incubation period, the mice were returned to the operant chamber to undergo a post-abstinence seeking test (PA) during which the active nosepoke resulted in the presentation of the contingent cues associated with the reward, but the sucrose reward was not delivered. After PA testing, mice underwent extinction training to suppress responding to contextual cues. Each extinction session lasted 120 mins (preceded by the 15 mins “light-OFF” period) during which the active nosepoke no longer resulted in the presentation of cues or delivery of the reward. Extinction criteria was determined *a priori* as two consecutive sessions with an average of presses less than 10 nosepokes. At least five extinction training sessions were conducted before the mice underwent reinstatement to the cues. Once the extinction criteria were met, mice underwent a cued-seeking reinstatement test (RST) for 2 hours during which the conditioned cues were returned after every active nosepoke. At the beginning of the reinstatement session, all mice received one free cue at the beginning of the session to elicit operant responding.

##### Cocaine Self-administration

After recovery from catheterization surgery, mice were trained daily (1 session/day) to nosepoke the active port to receive a cocaine injection (0.75mg/kg/infusion), which was associated with the presentation of a complex cue (light in the active port and a sound) for 5 seconds, followed by a 20-second time-out period, during which active nosepokes elicited no response. A maximum of 100 cocaine injections per session was allowed to prevent an overdose. Each session started with a 15-minute “baseline” period during which the house light was off and the nosepokes did not result in sucrose delivery, followed by 120-minutes of cocaine self-administration. The “baseline” period was used to collect a baseline recording during the calcium imaging experiments, or to adapt the animal to the headcap connections.

##### Post-Abstinence Seeking Test, Extinction, and Cued Reinstatement

After self-administration, mice undergo abstinence from cocaine self-administration for 7-10 days in their home cage during which they were handled daily. After the incubation period, the mice were returned to the operant chamber to undergo a post-abstinence seeking test (PA) for 120 minutes (preceded by the 15 mins “light-OFF” period) during which the active nosepoke resulted in the presentation of the contingent cues associated with the reward, but the cocaine reward was not delivered. For experiments including optogenetic and chemogenetic manipulations, multiple PA seeking tests were performed to assess persistence of the behavioral responses. After PA testing, mice underwent extinction training to suppress responding to contextual cues. Each extinction session lasted 120 mins (preceded by the 15 mins “light-OFF” period) during which the active nosepoke no longer resulted in the presentation of cues or delivery of the reward. Extinction criterion was determined *a priori* as two consecutive sessions with an average less than 10 nosepokes. At least five extinction training sessions were conducted before the mice underwent reinstatement to the cues. Once the extinction criterion was met, mice underwent a cued-seeking reinstatement test (RST) for 2 hours during which the conditioned cues were returned after every active nosepoke. At the beginning of the reinstatement session, all mice received one free cue at the beginning of the session to elicit operant responding. Mice were considered successfully reinstated in response to the cue if they nosepoke for the cue >10 times during a single session. Mice that failed a reinstatement test were given 1 more day of extinction before repeating the reinstatement test. Mice that failed two cued-reinstatement tests were excluded from the reinstatement test analysis.

#### Calcium Imaging from freely behaving animals

##### Acquisition

We recorded calcium activity of D1- and D2-MSN in freely behaving D1- and D2/A2A-cre mice respectively using miniature fluorescent microscopes “miniscopes” (nVista 3.0, Inscopix) on intermittent non-consecutive days throughout self-administration, post-abstinence seeking test, extinction training and reinstatement sessions. After the lens placement surgery, calcium dynamics were monitored weekly until individual neuron calcium changes were detected. These sessions were also used to adapt the mice for the weight of the miniscope. These mice were then randomized into cocaine self-administration and sucrose self-administration groups. Before the beginning of the first recording session, the gain (limit 2-4), the LED intensity (0.5-1 mW/mm^2^), and the desired z-plane were optimized per animal to obtain the highest number of well-defined regions of interests (ROIs), or putative neurons. The gain and LED settings were saved to be used in all subsequent sessions; the z-plane was electronically adjusted to match the same field of view during every session to allow tracking of the same neurons across time. All recordings were performed at 15 Hz (exposure time = 0.667 seconds), at 2x spatial down-sampling rate to reduce the generated file size. The operant chambers were connected to the Inscopix Data Acquisition Box using two transistor-transistor logic (TTL) via BNC cables to synchronize the recorded calcium imaging frames and behavioral events.

#### Image Processing and Signal Extraction

Processing and ROI segmentation. The pre-processing and ROI segmentation were performed using the MATLAB application programming interface (API) of Inscopix Data Processing Software v1.6 or custom MATLAB codes. The generated recordings were spatially down-sampled (2x in both dimensions) to reduce the file size and allow shorter processing times. Any defective pixels or dropped frames were corrected by linear interpolation from nearby pixels and frames. A spatial bandpass filer (0.005-0.5 pixels-1) was next applied to remove the low and high spatial frequency components that do not correspond to cells in focus. This was followed by applying a frame-by-frame motion correction algorithm to account for any motion artifacts. The signal was then normalized to background by subtracting and dividing each pixel value by the mean intensity of that pixel over the entire recording, as follow:

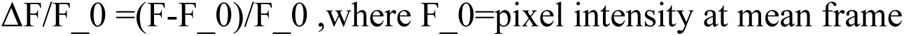

A principal component analysis (PCA), followed by an independent component analysis (ICA), were used to segment the recording into independent spatial footprints of putative neurons. The corresponding temporal trace of each spatial footprint was calculated by applying the spatial footprint on the background normalized movie. All components were manually inspected and included in the analysis if they show typical neuronal shape of the spatial footprints and canonical calcium signals of their corresponding temporal trace. If two neighboring neurons (distance<5 pixels) have highly correlated temporal traces (coefficient>0.9), the neuron with the lower signal- to-noise ratio (SNR) is excluded from the analysis to eliminate any duplicate components belonging to the same neuron.

Detection of Excitatory Calcium Events. Ca2+ excitatory events were detected using Inscopix Data Processing Software whenever the amplitude of the temporal trace crossed a 6 median absolute deviation (MAD), with an indicator decay time of 200ms. The time of the Ca2+ event occurrence was determined as the start of the event.

#### Histology

At the end of the behavioral experiments, the animals were deeply anesthetized with isoflurane and then transcardially perfused with 10mL of phosphate buffer-saline (PBS) followed by 20mL of 10% formalin solution. The brains were extracted and fixed in 10% formalin solution overnight and then transferred to 20% sucrose in PBS for cryoprotection. After 24 hours, the brains were frozen in dry ice and cut into 40μm sections using a cryostat. For immunohistochemical staining, free-floating sections were blocked with 2% normal donkey serum in 2% PBST for 1 hour on a shaker at room temperature. These sections were incubated overnight at 4⁰C in blocking solution containing chicken anti-GFP antibody (1:1000, Abcam ab13970). The next day, the sections were washed 3x for 5-minutes with 0.1% PBST solution, before staining them with standard Alexa conjugated secondary antibody (488 Goat Anti-Chicken, Thermofischer scientific: A-11039) for 2 hours. Hoechst 33258 (1:2000, Thermofischer) was added to slides during the last 15 minutes of incubation with secondary antibody to stain for nuclei. Finally, brain sections were washed, mounted on glass slides, and cover-slipped after adding Prolong Fold Fluorescent mounting medium. To ensure successful virus expression and accurate lens placement, confirmation was conducted following the staining under fluorescent microscopy.

### Quantification and statistical analysis

#### Calcium Imaging Analysis

##### Time-locked neurons

Time-locked neurons were defined similarly to what was previously done^54^. Briefly, the actual calcium activity of all neurons around the active nosepokes (−5s before to 10s after) was extracted, aligned to nosepoke onset, and averaged over the number of trials of interest (all or first 10 nosepokes). For every neuron, the Ca^2+^ activity around each rewarded nosepoke was normalized to a baseline taken between −5 to −3 seconds before the nosepoke. The averaged 15-second fluorescent neurons across the first 10 nosepokes were used to plot the peri-stimulus heatmaps for the represented sessions. The average trace of all neurons was compared to a 1000x shuffled distribution of means, obtained by circularly shuffling the calcium trace with respect to the nosepoke timestamps 1000x and generating a null distribution of peri-stimulus histograms. 95% confidence intervals of the null distribution are plotted to compare to the trace generated from real data.

To test whether a neuron was time-locked inhibited or time-locked excited, the maximum and minimum mean activity of each neuron was compared to a shuffled distribution of 1000 maxima and 1000 minima expected by chance. The shuffled distribution was generated by circularly shuffling the temporal trace of every neuron around the behavioral events by random values and calculating the minimum and maximum random mean activity at every iteration. A neuron was considered significantly inhibited if its actual minimum was lower than the 2.5^th^ percentile of the shuffled distribution of minima, and significantly excited if its actual maximum was higher than the 97.5^th^ percentile of the shuffled distribution of maxima.

The stability of the time-locked neurons within the same session was assessed by using odd-numbered events and even-numbered events. The neurons determined to be time-locked with similar activity patterns (excited or inhibited) to both sets were considered stable. All events were used in these analyses. Second, we repeated the time-locked analysis using the first 10 rewarded nosepokes and the second 10 rewarded nosepokes. The neurons determined to be time-locked with similar activity patterns to both sets were considered stable.

##### Tracking neurons across different sessions

Neurons between multiple sessions were tracked using a previously established nearest-neighbor cell registration method^19,55^. Maximum intensity projections for each session were generated and thresholded at 50% of the maximum pixel value, and the generated FOVs between sessions were first aligned to a reference session (last session of self-administration), using translational corrections only. A threshold for a candidate cell to be used for registration was determined by calculating within-session distances between nearest neighbors. Nearest neighbor centroid distances were found to be always greater than 4µm; we chose 3μm as a threshold to further reduce the chance of incorrect cross-session registration. Neurons were tracked between two late self-administration sessions. Neurons were also tracked between either of the two recorded self-administration sessions and the post-abstinence session: the pairing resulting in a higher number of tracked neurons was selected for analysis to maximize the power of the analyses.

##### Decoding Analysis

The nosepokes and cues were decoded using support vector machine models trained on the neural data 5 second before or 5 second after the nosepoke occurrence, respectively. For every event, a matched non-event epoch was randomly selected from the session, such as it is 30 seconds separated from any behavioral event in the session. Four-fold cross-validation was used to evaluate the ability of the model to decode the validation set. After the four iterations, a mean decoding accuracy is calculated and used to assess the ability of the neuronal data to predict the behavioral events. The labels of the events and non-events were shuffled to generate the shuffled distribution. For ensemble decoding, the ensembles were selected based on time-locked activity across all events. All decoding accuracy scores were compared between animals using 2-way ANOVA, followed by Bonferroni correction. For decoding analysis in longitudinally tracked neurons, an SVM model was trained using all the data from one session (SA), the resultant model was then applied to a test data from the test session (SA or PA). The performance of all models were compared to models trained on neuronal with shuffled event labels. Data was plotted and reported as decoding accuracy using the real data subtracted from the decoding accuracy when using the shuffled dataset.

#### Statistical Analysis and Graphics

Heatmaps and example traces were generated using MATLAB. Summary graphical representations were generated using GraphPad Prism v9. All statistical analysis were performed using either MATLAB 2022b or GraphPad Prism v9. All statistical rests were corrected for multiple comparisons with the Bonferroni method when applicable. Comparisons of fractions of neurons were done using Chi-Square tests. For the decoding results, all comparisons were made using paired Student’s t-tests (real vs shuffled) and p values were corrected using a false discovery rate with q=0.02. Animals in which there was <10 neurons were excluded from the analyses but included in the behavioral data in figure 1. For all analysis, a p-value of 0.05 was considered significant a priori.

## REFERENCES

1. Pennartz, C.M.A., Groenewegen, H.J., and Lopes da Silva, F.H. (1994). The nucleus accumbens as a complex of functionally distinct neuronal ensembles: An integration of behavioural, electrophysiological and anatomical data. Progress in Neurobiology 42, 719–761. 10.1016/0301-0082(94)90025-6.

2. Bobadilla, A.-C., Heinsbroek, J.A., Gipson, C.D., Griffin, W.C., Fowler, C.D., Kenny, P.J., and Kalivas, P.W. (2017). Chapter 4 - Corticostriatal plasticity, neuronal ensembles, and regulation of drug-seeking behavior. In Progress in Brain Research, T. Calvey, and W.M.U. Daniels, eds. (Elsevier), pp. 93–112. 10.1016/bs.pbr.2017.07.013.

3. Cruz, F.C., Koya, E., Guez-Barber, D.H., Bossert, J.M., Lupica, C.R., Shaham, Y., and Hope, B.T. (2013). New technologies for examining the role of neuronal ensembles in drug addiction and fear. Nature Reviews Neuroscience 14, 743–754. 10.1038/nrn3597.

4. Bobadilla, A.-C., Dereschewitz, E., Vaccaro, L., Heinsbroek, J.A., Scofield, M.D., and Kalivas, P.W. (2020). Cocaine and sucrose rewards recruit different seeking ensembles in the nucleus accumbens core. Molecular Psychiatry 25, 3150–3163. 10.1038/s41380-020-00888-z.

5. Carelli, R.M. (2002). Nucleus accumbens cell firing during goal-directed behaviors for cocaine vs. ‘natural’ reinforcement. Physiology & behavior 76, 379–387. 10.1016/S0031-9384(02)00760-6.

6. Roitman, M.F., Wheeler, R.A., and Carelli, R.M. (2005). Nucleus accumbens neurons are innately tuned for rewarding and aversive taste stimuli, encode their predictors, and are linked to motor output. Neuron 45, 587–597. 10.1016/j.neuron.2004.12.055.

7. Van Zessen, R., Li, Y., Marion-Poll, L., Hulo, N., Flakowski, J., and Lüscher, C. (2021). Dynamic dichotomy of accumbal population activity underlies cocaine sensitization. eLife 10. 10.7554/elife.66048.

8. Grant, R.I., Doncheck, E.M., Vollmer, K.M., Winston, K.T., Romanova, E.V., Siegler, P.N., Holman, H., Bowen, C.W., and Otis, J.M. (2021). Specialized coding patterns among dorsomedial prefrontal neuronal ensembles predict conditioned reward seeking. eLife 10. 10.7554/elife.65764.

9. Taniguchi, M., Carreira, M.B., Cooper, Y.A., Bobadilla, A.C., Heinsbroek, J.A., Koike, N., Larson, E.B., Balmuth, E.A., Hughes, B.W., Penrod, R.D., et al. (2017). HDAC5 and Its Target Gene, Npas4, Function in the Nucleus Accumbens to Regulate Cocaine-Conditioned Behaviors. Neuron 96, 130–144.e136. 10.1016/j.neuron.2017.09.015.

10. Sortman, B.W., Rakela, S., Paprotna, S., Cerci, B., and Warren, B.L. (2024). Nucleus accumbens neuronal ensembles vary with cocaine reinforcement in male and female rats. Addict Biol 29, e13397. 10.1111/adb.13397.

11. Scofield, M.D., Heinsbroek, J.A., Gipson, C.D., Kupchik, Y.M., Spencer, S., Smith, A.C.W., Roberts-Wolfe, D., and Kalivas, P.W. (2016). The Nucleus Accumbens: Mechanisms of Addiction across Drug Classes Reflect the Importance of Glutamate Homeostasis. Pharmacological Reviews 68, 816–871. 10.1124/pr.116.012484.

12. Carelli, R.M., and Wondolowski, J. (2003). Selective Encoding of Cocaine versus Natural Rewards by Nucleus Accumbens Neurons Is Not Related to Chronic Drug Exposure. The Journal of Neuroscience 23, 11214–11223. 10.1523/jneurosci.23-35-11214.2003.

13. Namboodiri, V.M.K., Otis, J.M., van Heeswijk, K., Voets, E.S., Alghorazi, R.A., Rodriguez-Romaguera, J., Mihalas, S., and Stuber, G.D. (2019). Single-cell activity tracking reveals that orbitofrontal neurons acquire and maintain a long-term memory to guide behavioral adaptation. Nature Neuroscience 22, 1110–1121. 10.1038/s41593-019-0408-1.

14. Resendez, S.L., Jennings, J.H., Ung, R.L., Namboodiri, V.M.K., Zhou, Z.C., Otis, J.M., Nomura, H., McHenry, J.A., Kosyk, O., and Stuber, G.D. (2016). Visualization of cortical, subcortical and deep brain neural circuit dynamics during naturalistic mammalian behavior with head-mounted microscopes and chronically implanted lenses. Nature Protocols 11, 566. 10.1038/nprot.2016.021.

15. Tunstall, B.J., and Kearns, D.N. (2016). Cocaine can generate a stronger conditioned reinforcer than food despite being a weaker primary reinforcer. Addict Biol 21, 282–293. 10.1111/adb.12195.

16. Wise, R.A., and Kiyatkin, E.A. (2011). Differentiating the rapid actions of cocaine. Nature Reviews Neuroscience 12, 479–484. 10.1038/nrn3043.

17. Surmeier, D.J., Ding, J., Day, M., Wang, Z., and Shen, W. (2007). D1 and D2 dopamine-receptor modulation of striatal glutamatergic signaling in striatal medium spiny neurons. Trends in neurosciences 30, 228–235. S0166-2236(07)00069-0 [pii] 10.1016/j.tins.2007.03.008.

18. Canchy, L., Girardeau, P., Durand, A., Vouillac-Mendoza, C., and Ahmed, S.H. (2021). Pharmacokinetics trumps pharmacodynamics during cocaine choice: a reconciliation with the dopamine hypothesis of addiction. Neuropsychopharmacology 46, 288–296. 10.1038/s41386-020-0786-9.

19. Sheintuch, L., Rubin, A., Brande-Eilat, N., Geva, N., Sadeh, N., Pinchasof, O., and Ziv, Y. (2017). Tracking the Same Neurons across Multiple Days in Ca2+ Imaging Data. Cell Reports 21, 1102–1115. 10.1016/j.celrep.2017.10.013.

20. Stuber, G.D., Roitman, M.F., Phillips, P.E.M., Carelli, R.M., and Wightman, R.M. (2005). Rapid Dopamine Signaling in the Nucleus Accumbens during Contingent and Noncontingent Cocaine Administration. Neuropsychopharmacology 30, 853–863. 10.1038/sj.npp.1300619.

21. Lee, R.S., Koob, G.F., and Henriksen, S.J. (1998). Electrophysiological responses of nucleus accumbens neurons to novelty stimuli and exploratory behavior in the awake, unrestrained rat. Brain Res 799, 317–322. 10.1016/s0006-8993(98)00477-6.

22. Eoin, Kremer, Y., Lefort, S., Harada, M., Pascoli, V., Rohner, C., and Lüscher, C. (2015). Accumbal D1R Neurons Projecting to Lateral Hypothalamus Authorize Feeding. Neuron 88, 553–564. 10.1016/j.neuron.2015.09.038.

23. Smith, R.J., Lobo, M.K., Spencer, S., and Kalivas, P.W. (2013). Cocaine-induced adaptations in D1 and D2 accumbens projection neurons (a dichotomy not necessarily synonymous with direct and indirect pathways). Curr Opin Neurobiol 23, 546–552. 10.1016/j.conb.2013.01.026.

24. Anderson, E.M., and Self, D.W. (2017). It’s only a matter of time: longevity of cocaine-induced changes in dendritic spine density in the nucleus accumbens. Curr Opin Behav Sci 13, 117–123. 10.1016/j.cobeha.2016.11.013.

25. Kim, J., Park, B.H., Lee, J.H., Park, S.K., and Kim, J.H. (2011). Cell type-specific alterations in the nucleus accumbens by repeated exposures to cocaine. Biological Psychiatry 69, 1026–1034. 10.1016/j.biopsych.2011.01.013.

26. Ma, Y.Y., Lee, B.R., Wang, X., Guo, C., Liu, L., Cui, R., Lan, Y., Balcita-Pedicino, J.J., Wolf, M.E., Sesack, S.R., et al. (2014). Bidirectional modulation of incubation of cocaine craving by silent synapse-based remodeling of prefrontal cortex to accumbens projections. Neuron 83, 1453–1467. 10.1016/j.neuron.2014.08.023.

27. Conrad, K.L., Tseng, K.Y., Uejima, J.L., Reimers, J.M., Heng, L.J., Shaham, Y., Marinelli, M., and Wolf, M.E. (2008). Formation of accumbens GluR2-lacking AMPA receptors mediates incubation of cocaine craving. Nature 454, 118–121. 10.1038/nature06995.

28. Anderson, S.M., Famous, K.R., Sadri-Vakili, G., Kumaresan, V., Schmidt, H.D., Bass, C.E., Terwilliger, E.F., Cha, J.H., and Pierce, R.C. (2008). CaMKII: a biochemical bridge linking accumbens dopamine and glutamate systems in cocaine seeking. Nat Neurosci 11, 344–353. 10.1038/nn2054.

29. Pascoli, V., Terrier, J., Espallergues, J., Valjent, E., O’Connor, E.C., and Lüscher, C. (2014). Contrasting forms of cocaine-evoked plasticity control components of relapse. Nature 509, 459-464. 10.1038/nature13257.

30. Roberts-Wolfe, D., Bobadilla, A.-C., Heinsbroek, J.A., Neuhofer, D., and Kalivas, P.W. (2018). Drug Refraining and Seeking Potentiate Synapses on Distinct Populations of Accumbens Medium Spiny Neurons. The Journal of Neuroscience 38, 7100–7107. 10.1523/jneurosci.0791-18.2018.

31. Heinsbroek, J.A., Neuhofer, D.N., Griffin, W.C., Siegel, G.S., Bobadilla, A.-C., Kupchik, Y.M., and Kalivas, P.W. (2017). Loss of Plasticity in the D2-Accumbens Pallidal Pathway Promotes Cocaine Seeking. The Journal of Neuroscience 37, 757–767. 10.1523/jneurosci.2659-16.2016.

32. Garcia-Keller, C., Scofield, M.D., Neuhofer, D., Varanasi, S., Reeves, M.T., Hughes, B., Anderson, E., Richie, C.T., Mejias-Aponte, C., Pickel, J., et al. (2020). Relapse-Associated Transient Synaptic Potentiation Requires Integrin-Mediated Activation of Focal Adhesion Kinase and Cofilin in D1-Expressing Neurons. The Journal of neuroscience : the official journal of the Society for Neuroscience 40, 8463–8477. 10.1523/JNEUROSCI.2666-19.2020.

33. Chioma, V.C., Kruyer, A., Bobadilla, A.C., Angelis, A., Ellison, Z., Hodebourg, R., Scofield, M.D., and Kalivas, P.W. (2021). Heroin Seeking and Extinction From Seeking Activate Matrix Metalloproteinases at Synapses on Distinct Subpopulations of Accumbens Cells. Biol Psychiatry 89, 947–958. 10.1016/j.biopsych.2020.12.004.

34. Garcia-Keller, C., Neuhofer, D., Bobadilla, A.C., Spencer, S., Chioma, V.C., Monforton, C., and Kalivas, P.W. (2019). Extracellular Matrix Signaling Through beta3 Integrin Mediates Cocaine Cue-Induced Transient Synaptic Plasticity and Relapse. Biol Psychiatry 86, 377–387. 10.1016/j.biopsych.2019.03.982.

35. Kruyer, A., and Kalivas, P.W. (2021). Astrocytes as cellular mediators of cue reactivity in addiction. Curr Opin Pharmacol 56, 1–6. 10.1016/j.coph.2020.07.009.

36. Lobo, M.K., Covington, H.E., Chaudhury, D., Friedman, A.K., Sun, H., Damez-Werno, D., Dietz, D.M., Zaman, S., Koo, J.W., Kennedy, P.J., et al. (2010). Cell Type–Specific Loss of BDNF Signaling Mimics Optogenetic Control of Cocaine Reward. Science 330, 385–390. 10.1126/science.1188472.

37. Pardo-Garcia, T.R., Garcia-Keller, C., Penaloza, T., Richie, C.T., Pickel, J., Hope, B.T., Harvey, B.K., Kalivas, P.W., and Heinsbroek, J.A. (2019). Ventral Pallidum Is the Primary Target for Accumbens D1 Projections Driving Cocaine Seeking. The Journal of Neuroscience 39, 2041–2051. 10.1523/jneurosci.2822-18.2018.

38. Calipari, E.S., Bagot, R.C., Purushothaman, I., Davidson, T.J., Yorgason, J.T., Peña, C.J., Walker, D.M., Pirpinias, S.T., Guise, K.G., Ramakrishnan, C., et al. (2016). In vivo imaging identifies temporal signature of D1 and D2 medium spiny neurons in cocaine reward. Proceedings of the National Academy of Sciences 113, 2726–2731. 10.1073/pnas.1521238113.

39. Bock, R., Shin, J.H., Kaplan, A.R., Dobi, A., Markey, E., Kramer, P.F., Gremel, C.M., Christensen, C.H., Adrover, M.F., and Alvarez, V.A. (2013). Strengthening the accumbal indirect pathway promotes resilience to compulsive cocaine use. Nat Neurosci 16, 632–638. 10.1038/nn.3369.

40. Soares-Cunha, C., Coimbra, B., David-Pereira, A., Borges, S., Pinto, L., Costa, P., Sousa, N., and Rodrigues, A.J. (2016). Activation of D2 dopamine receptor-expressing neurons in the nucleus accumbens increases motivation. Nature Communications 7, 11829. 10.1038/ncomms11829.

41. Soares-Cunha, C., Coimbra, B., Sousa, N., and Rodrigues, A.J. (2016). Reappraising striatal D1- and D2-neurons in reward and aversion. Neuroscience & Biobehavioral Reviews 68, 370–386. 10.1016/j.neubiorev.2016.05.021.

42. Soares-Cunha, C., Coimbra, B., Domingues, A.V., Vasconcelos, N., Sousa, N., and Rodrigues, A.J. (2018). Nucleus Accumbens Microcircuit Underlying D2-MSN-Driven Increase in Motivation. eNeuro 5. 10.1523/eneuro.0386-18.2018.

43. Lafferty, C.K., Yang, A.K., Mendoza, J.A., and Britt, J.P. (2020). Nucleus Accumbens Cell Type- and Input-Specific Suppression of Unproductive Reward Seeking. Cell Reports 30, 3729–3742.e3723. 10.1016/j.celrep.2020.02.095.

44. Pedersen, C.E., Castro, D.C., Gray, M.M., Zhou, Z.C., Piantadosi, S.C., Gowrishankar, R., Kan, S.A., Murphy, P.J., O’Neill, P.R., and Bruchas, M.R. (2022). Medial accumbens shell spiny projection neurons encode relative reward preference. Cold Spring Harbor Laboratory.

45. Zachry, J.E., Kutlu, M.G., Yoon, H.J., Leonard, M.Z., Chevée, M., Patel, D.D., Gaidici, A., Kondev, V., Thibeault, K.C., Bethi, R., et al. (2024). D1 and D2 medium spiny neurons in the nucleus accumbens core have distinct and valence-independent roles in learning. Neuron 112, 835–849.e837. 10.1016/j.neuron.2023.11.023.

46. Zhao, Z.D., Han, X., Chen, R., Liu, Y., Bhattacherjee, A., Chen, W., and Zhang, Y. (2022). A molecularly defined D1 medium spiny neuron subtype negatively regulates cocaine addiction. Sci Adv 8, eabn3552. 10.1126/sciadv.abn3552.

47. McGlinchey, E.M., James, M.H., Mahler, S.V., Pantazis, C., and Aston-Jones, G. (2016). Prelimbic to Accumbens Core Pathway Is Recruited in a Dopamine-Dependent Manner to Drive Cued Reinstatement of Cocaine Seeking. The Journal of Neuroscience 36, 8700–8711. 10.1523/jneurosci.1291-15.2016.

48. Nall, R.W., Heinsbroek, J.A., Nentwig, T.B., Kalivas, P.W., and Bobadilla, A.C. (2021). Circuit selectivity in drug versus natural reward seeking behaviors. Journal of neurochemistry 157, 1450–1472. 10.1111/jnc.15297.

49. Pascoli, V., Turiault, M., and Lüscher, C. (2012). Reversal of cocaine-evoked synaptic potentiation resets drug-induced adaptive behaviour. Nature 481, 71–75. 10.1038/nature10709.

50. Saunders, B.T., Yager, L.M., and Robinson, T.E. (2013). Cue-evoked cocaine “craving”: role of dopamine in the accumbens core. The Journal of neuroscience : the official journal of the Society for Neuroscience 33, 13989–14000. 10.1523/jneurosci.0450-13.2013.

51. Gipson, Cassandra D., Kupchik, Yonatan M., Shen, H., Reissner, Kathryn J., Thomas, Charles A., and Kalivas, Peter W. (2013). Relapse Induced by Cues Predicting Cocaine Depends on Rapid, Transient Synaptic Potentiation. Neuron 77, 867–872. 10.1016/j.neuron.2013.01.005.

52. Heinsbroek, J.A., Bobadilla, A.C., Dereschewitz, E., Assali, A., Chalhoub, R.M., Cowan, C.W., and Kalivas, P.W. (2020). Opposing Regulation of Cocaine Seeking by Glutamate and GABA Neurons in the Ventral Pallidum. Cell Rep 30, 2018–2027.e2013. 10.1016/j.celrep.2020.01.023.

53. Kmiotek, E.K., Baimel, C., and Gill, K.J. (2012). Methods for Intravenous Self Administration in a Mouse Model. Journal of Visualized Experiments. 10.3791/3739.

54. Cameron, C.M., Murugan, M., Choi, J.Y., Engel, E.A., and Witten, I.B. (2019). Increased Cocaine Motivation Is Associated with Degraded Spatial and Temporal Representations in IL-NAc Neurons. Neuron 103, 80–91.e87. 10.1016/j.neuron.2019.04.015.

55. Rubin, A., Geva, N., Sheintuch, L., and Ziv, Y. (2015). Hippocampal ensemble dynamics timestamp events in long-term memory. eLife 4, e12247. 10.7554/eLife.12247.

