## Supplemental Tables for "Formation of an Enduring Ensemble of Accumbens Neurons Leads to Prepotent Seeking for Cocaine Over Natural Reward Cues"

**Table 1.** Chi^2^ values comparing proportion of stable neurons in the MSN excited (Ex) or inhibited (In) subpopulations between sucrose and cocaine in the MSN excited and inhibited subpopulations using the odd/even analysis of stability. Statistics correspond to Figure 4b-d.

|  | D1 Ex | D1 In | D2 Ex | D2 In |
| --- | --- | --- | --- | --- |
| SA | 3.577, p= 0.059 | 2.616, p= 0.128 | 2.331, p= 0.137 | 1.010, p= 0.330 |
| PA | **4.164, p= 0.048** | 1.326, p= 0.316 | 0.008, p> 0.999 | **8.996, p= 0.003** |
| RST | 3.343, p= 0.068 | 0.671, p= 0.665 | 1.456, p= 0.401 | 1.890, p= 0.263 |

Bold: statistically significant (p<0.05); Data shown as Chi^2^ values and p values comparing sucrose with cocaine within each subpopulation. (Ex: excited, In: inhibited)

**Table 2.** Correlation coefficients comparing subpopulation percentage to total cued lever presses across the 120 min self-administration (SA), post-abstinence (PA) and reinstatement (RST) sessions for sucrose- and cocaine-trained mice. R^2^ values correspond to Figure 4f.

|  | Excited Neurons | | | | | |
| --- | --- | --- | --- | --- | --- | --- |
|  | Sucrose | | | Cocaine | | |
|  | D1+D2 | D1 | D2 | D1+D2 | D1 | D2 |
| SA | 0.026 | 0.041 | 0.020 | **0.171** | **0.309** | 0.001 |
| PA | 0.031 | 0.202 | 0.238 | **0.383** | **0.724** | 0.040 |
| RST | 0.019 | 0.175 | 0.007 | 0.005 | <0.001 | 0.030 |

|  | Inhibited Neurons | | | | | |
| --- | --- | --- | --- | --- | --- | --- |
|  | Sucrose | | | Cocaine | | |
|  | D1+D2 | D1 | D2 | D1+D2 | D1 | D2 |
| SA | 0.023 | 0.175 | 0.003 | **0.384** | **0.416** | 0.003 |
| PA | 0.027 | <0.001 | <0.001 | 0.167 | 0.275 | 0.156 |
| RST | 0.055 | 0.028 | 0.147 | 0.118 | 0.249 | 0.044 |

Bold: statistically significant (p<0.05); Data shown as r^2^ values from a linear regression for each comparison. Correlating rewarded/cued (SA) or unrewarded/cued (PA; RST) NPs versus the percentage of each subpopulation. N= 6-7 for each comparison.

**Table 3.** Summary of Students t-test values and p values for decoding NPs in each subpopulation compared to the respective shuffled distribution. Statistics correspond to Figure 5.

| **SUCROSE** | D1 Ex | D1 In | D1 N | D2 Ex | D2 I | D2 N |
| --- | --- | --- | --- | --- | --- | --- |
| SA | **11.15, p<0.001** | 3.756, p= 0.009 | **5.537, p= 0.002** | **11.15, p< 0.001** | 3.756, p= 0.009 | **5.537, p= 0.002** |
| PA | **5.090, p= 0.004** | 4.430, p= 0.007 | 1.696, p= 0.151 | 3.252, p= 0.017 | 0.651, p= 0.539 | 1.023, p= 0.346 |
| RST | 2.217, p= 0.077 | 1.117, p= 0.293 | 1.593, p= 0.172 | 2.386, p= 0.027 | 0.049, p= 0.963 | 0.228, p= 0.827 |

| **COCAINE** | D1 Ex | D1 In | D1 N | D2 Ex | D2 I | D2 N |
| --- | --- | --- | --- | --- | --- | --- |
| SA | **4.495, p= 0.004** | **5.041, p= 0.002** | 2.362, p= 0.056 | 3.480, p= 0.013 | 3.540, p= 0.012 | 1.433, p= 0.202 |
| PA | **7.378, p< 0.001** | 4.702, p= 0.003 | 1.129, p= 0.302 | **5.982, p= 0.001** | 2.301, p= 0.061 | 0.581, p= 0.583 |
| RST | **4.848, p= 0.005** | 0.274, p= 0.795 | 1.125, p= 0.312 | 3.538, p= 0.012 | 3.000, p= 0.024 | 1.599, p= 0.165 |

Bold: statistically significant (p<0.05); using a paired Student’s t-test comparing real to shuffled data in each cell after adjustment using a false discovery rate, q= 0.02. Degree of freedom = 5-6 for each cell. (Ex: excited, In: inhibited)

**Table 4.** Chi^2^ values comparing proportion of stable neurons in the MSN subpopulations between sucrose and cocaine using the odd/even analysis of stability. Statistics correspond to Figure 6b-c, e-f.

|  | D1 Ex | D1 In | D2 Ex | D2 In |
| --- | --- | --- | --- | --- |
| SA to SA | 0.134, p= 0.824 | 0.726, p= 0.518 | 0.072, p= 0.808 | 0.602, p= 0.602 |
| SA to PA | **5.040, p= 0.029** | 1.978, p= 0.222 | 0.841, p= 0.401 | 0.024, p< 0.999 |

Bold: statistically significant (p<0.05); Data shown as Chi^2^ and p values from a 2 X 2 comparison (stable/unstable in sucrose/cocaine). Degrees of freedom = 1 for all comparisons. (Ex: excited, In: inhibited)

**Table 5.** Summary of Students t-test values and p values for decoding NPs in each subpopulation compared to the respective shuffled distribution, corresponding to data in Figure 6d,g).

|  | SUCROSE | | COCAINE | |
| --- | --- | --- | --- | --- |
|  | D1 MSN | D2 MSN | D1 MSN | D2 MSN |
| SA to SA | 5.175, p= 0.004 | 1.422, p= 0.214 | 4.137, p= 0.006 | 0.604, p= 0.568 |
| SA to PA | 0.646, p= 0.547 | 0.356, p= 0.386 | 2.634, p= 0.039 | 0.633, p= 0.555 |

Bold: statistically significant (p<0.05); using a paired Student’s t-test comparing quantified minus shuffled data between two self-administration sessions (SA-SA) or between a self-administration and post-abstinence session (SA to PA). Degrees of freedom = 5 or 6 in each cell. (Ex: excited, In: inhibited)
