## Supplemental Figures for "Formation of an Enduring Ensemble of Accumbens Neurons Leads to Prepotent Seeking for Cocaine Over Natural Reward Cues"

### Supplemental Figures and Legends

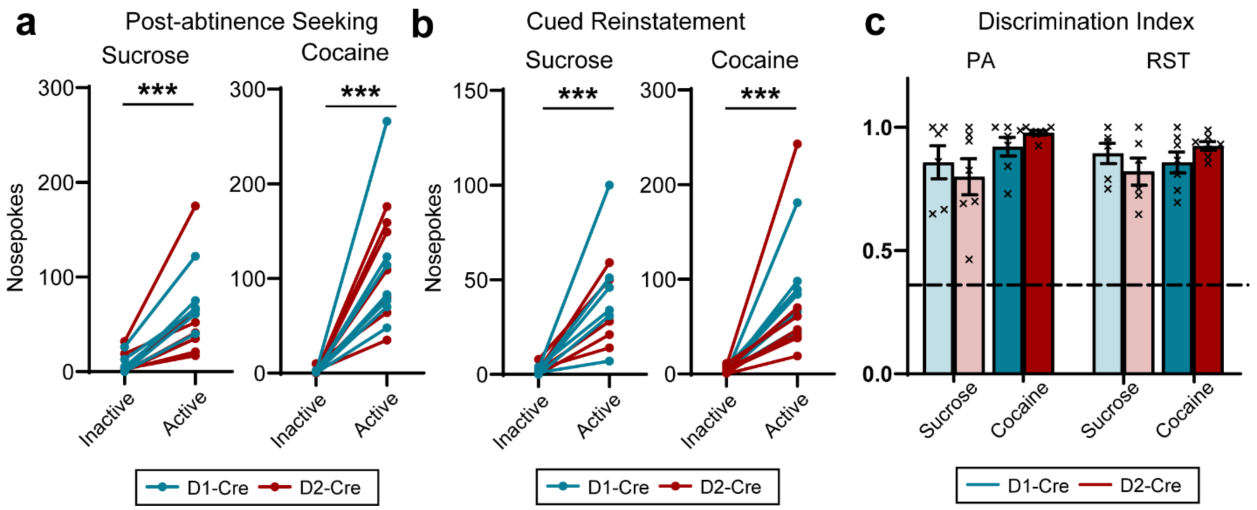


**Figure S1. D1- and D2-cre mice discriminate between active and inactive NPs during post-abstinence and cued reinstatement testing.**

**(a)** D1- and D2-cre mice pressed significantly more on active versus inactive NPs during sucrose (2-way ANOVA: NP x Genotype: $F_{(1,22)}= 0.319$, $p= 0.578$; NP: $F_{(1,22)}= 20.23$, $p<0.001$; Genotype: $F_{(1,22)}= 0.252$, $p= 0.620$) or cocaine (2-way ANOVA: NP x Genotype: $F_{(1,24)}= 0.007$, $p=0.934$; Nosepoke: $F_{(1,24)}= 40.93$, $p<0.001$; Genotype: $F_{(1,24)} <0.001$, $p= 0.993$) post-abstinence seeking. **(b)** D1- and D2-cre mice pressed significantly more on active versus inactive NPs during sucrose (2-way ANOVA: NP x Genotype: $F_{(1,20)}= 1.077$, $p= 0.312$; NP: $F_{(1,20)}= 21.15$, $p< 0.001$; Genotype: $F_{(1,20)}= 0.862$, $p= 0.364$) or cocaine (2-way ANOVA: NP x Genotype: $F_{(1,24} = 0.2012$, $p= 0.6578$; NP: $F_{(1,24)}= 20.58$, $p< 0.001$; Genotype: $F_{(1,24)}= 0.0981$, $p= 0.7569$) cued reinstatement. **(c)** Distribution of the discrimination index (calculated as, (active nosepoke - inactive nosepoke)/(active nosepoke + inactive nosepoke)) of D1- and D2-cre mice during post-abstinence and cued-reinstatement testing in sucrose and cocaine groups. Dotted line at 0.33 represents a 2:1 discrimination ratio.

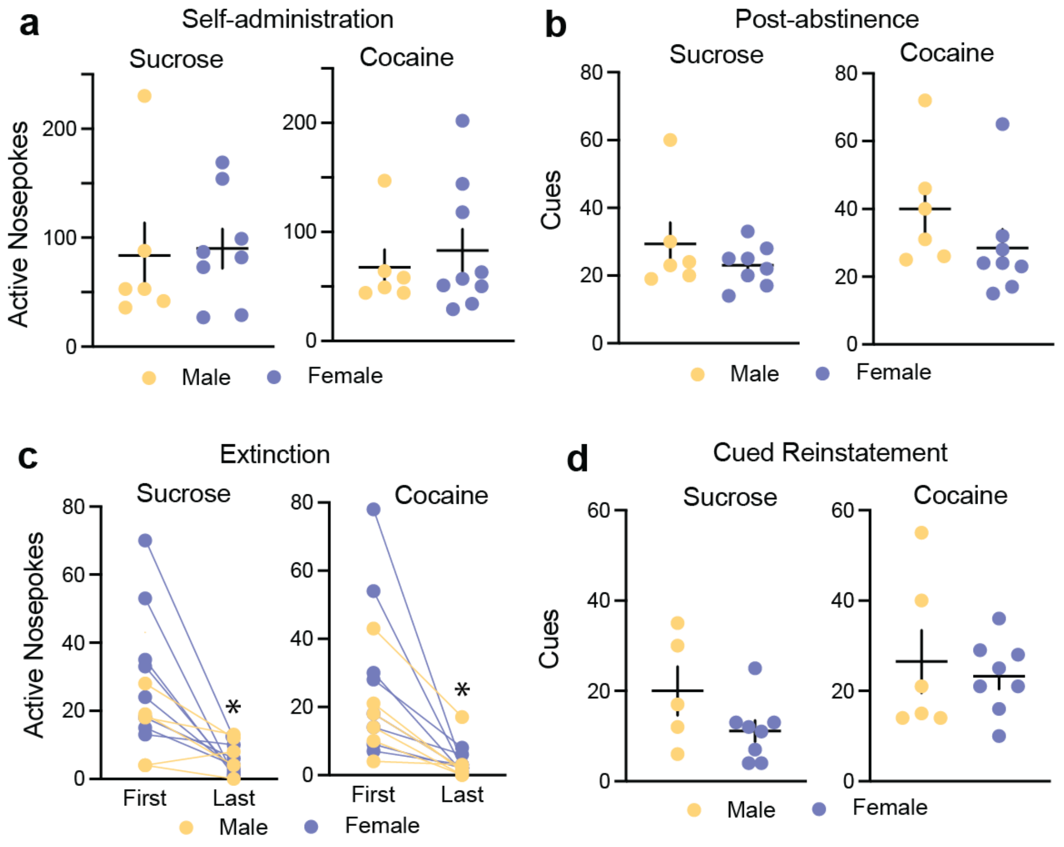


**Figure S2. Behavioral responding stratified by sex shows similar performance between male and female mice.** **(a)** Self-administration: Sucrose- unpaired $t_{(12)}= 0.190$, $p= 0.853$, Cocaine- unpaired $t_{(13)}= 0.561$, $p=0.584$. **(b)** Post-abstinence seeking: Sucrose- unpaired $t_{(12)}= 1.062$, $p= 0.309$, Cocaine- unpaired $t_{(13)}= 1/286$, $p=0.223$. **(c)** Extinction: Sucrose- 2-way RM-ANOVA: interaction: $F_{(1,11)}= 3.234$, $p= 0.100$; Sex: $F_{(1,12)}= 0.704$, $p= 0.418$; first vs last: $F_{(1,11)}= 16.91$, $p= 0.002$); Cocaine- 2-way RM-ANOVA: interaction: $F_{(1,12)}= 1.200$, $p= 0.291$; Sex: $F_{(1,12)}= 0.813$, $p= 0.385$; first vs last: $F_{(1,12)}= 14.45$, $p= 0.003$). *$p<0.05$ comparing first to last day of extinction. **(d)** Cued reinstatement: Sucrose- unpaired $t_{(11)}= 1.171$, $p= 0.116$, Cocaine- unpaired $t_{(12)}= 0.475$, $p=0.644$.

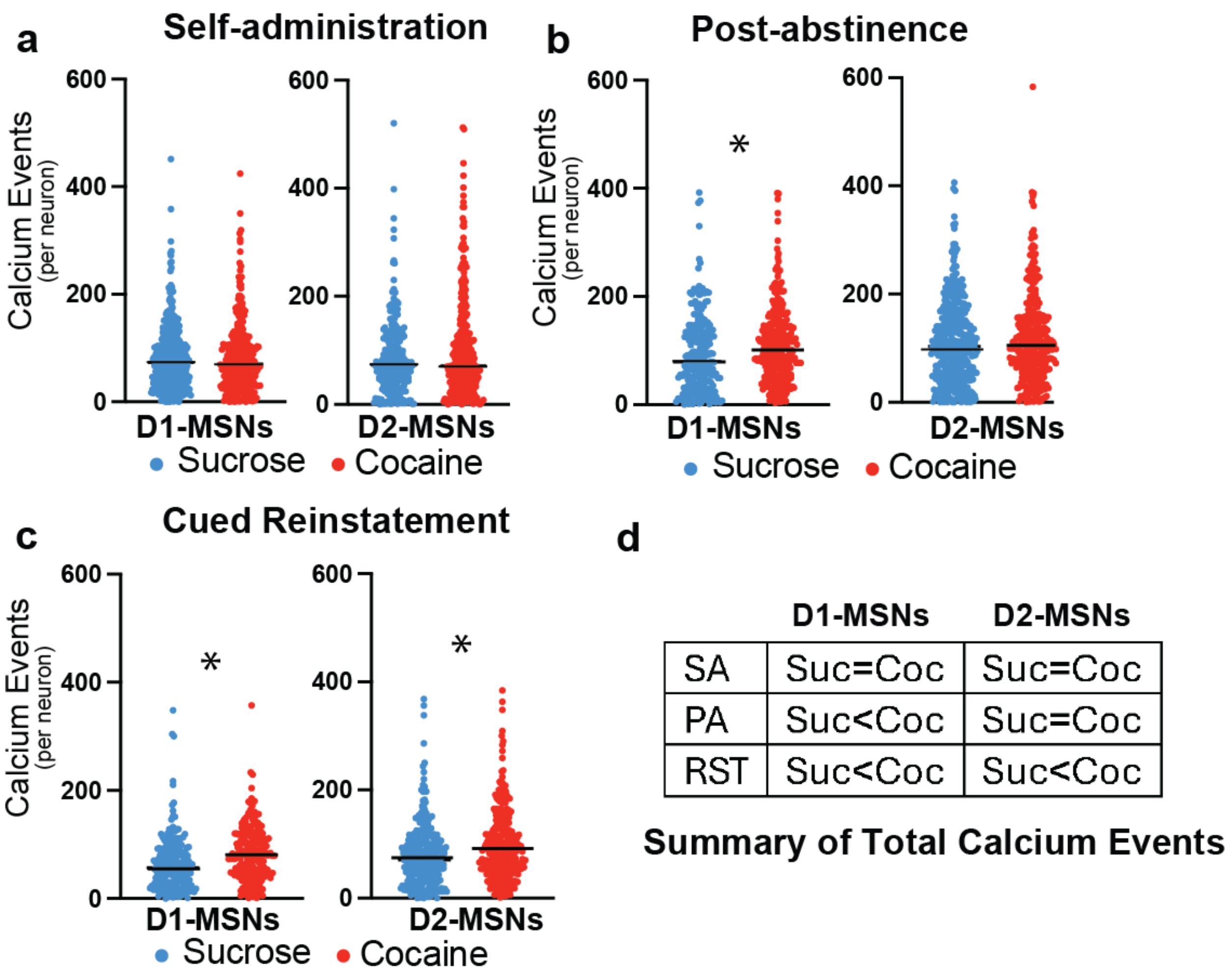


**Figure S3. D1-MSNs show an increase in total calcium events during unrewarded seeking in cocaine versus sucrose mice.** **(a)** Self-administration (SA): D1-MSNs- Mann-Whitney U= 55152, p= 0.685; D2-MSNs- Mann-Whitney U= 40616, p=0.769. **(b)** Post-abstinence (PA) seeking: D1-MSNs- Mann-Whitney U= 31013, p= 0.002, D2-MSNs- Mann-Whitney U= 56938, p=0.221. **(c)** Cue-reinstated (RST) seeking: D1-MSNs- Mann-Whitney U= 15872, p< 0.001, D2-MSNs- Mann-Whitney U= 38995, p=0.003. **(d)** Summary of number of $Ca^{2+}$ events showing that for unrewarded seeking, cocaine trained mice generally had more events than sucrose mice. *p< 0.05 comparing cocaine and sucrose.

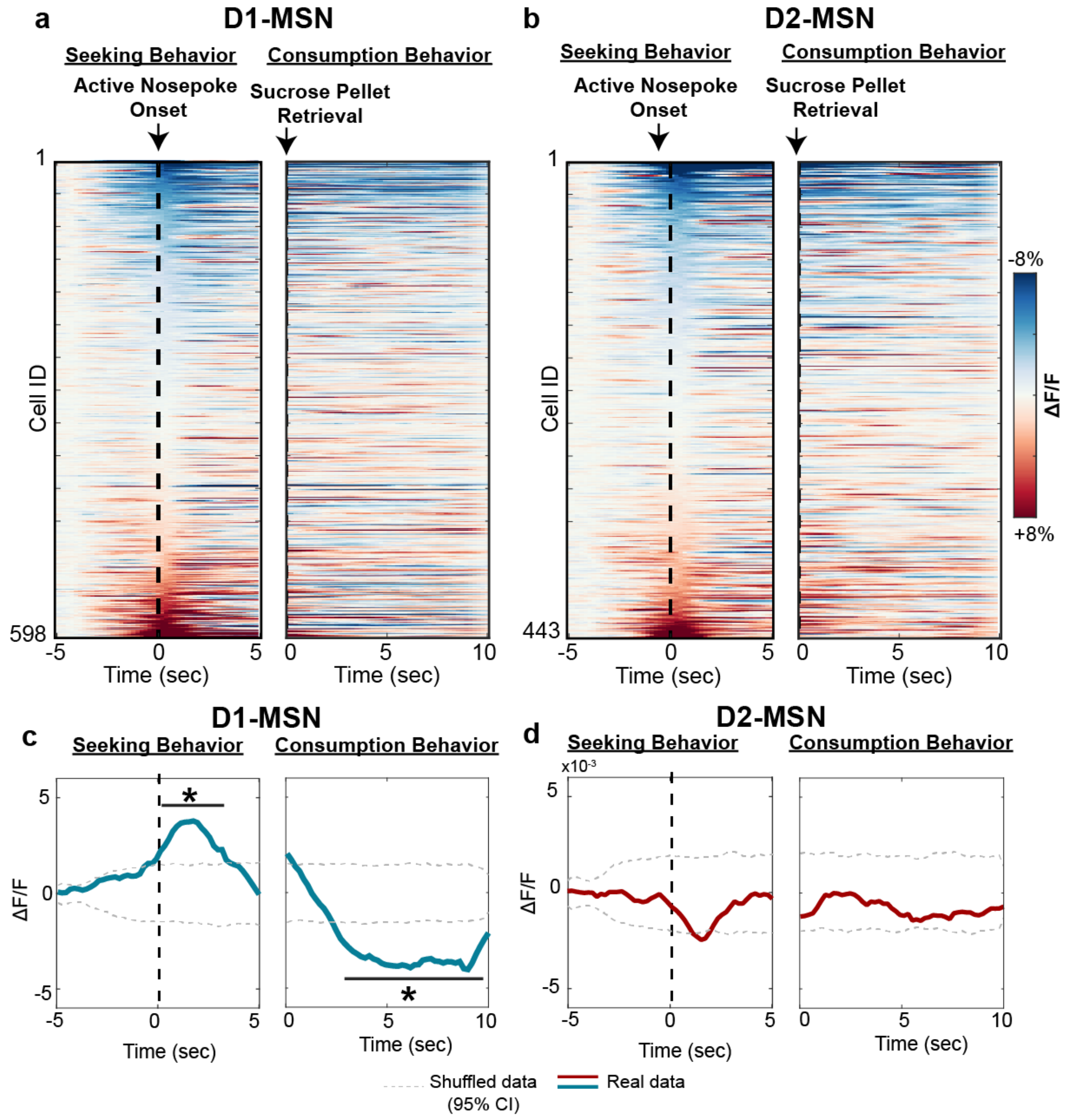


**Figure S4. Inhibitory population mean activity of D1-MSN during sucrose self-administration is timed to reward delivery.**

**(a,b)** Heatmaps representing the mean activity of recorded D1- and D2-MSN of the first 10 trials, aligned to the active nosepoke (left) and sucrose pellet consumption (right). **(c,d)** Mean activity traces of all recorded neurons (D1: blue, D2: red) compared to a shuffled distribution (Gray, dotted line represents 95% confidence interval, CI). D1-MSN show an excitatory activity following nosepoke, and inhibitory activity following sucrose consumption (compared to 95% CI from 1000x shuffled distribution, *p<0.05).

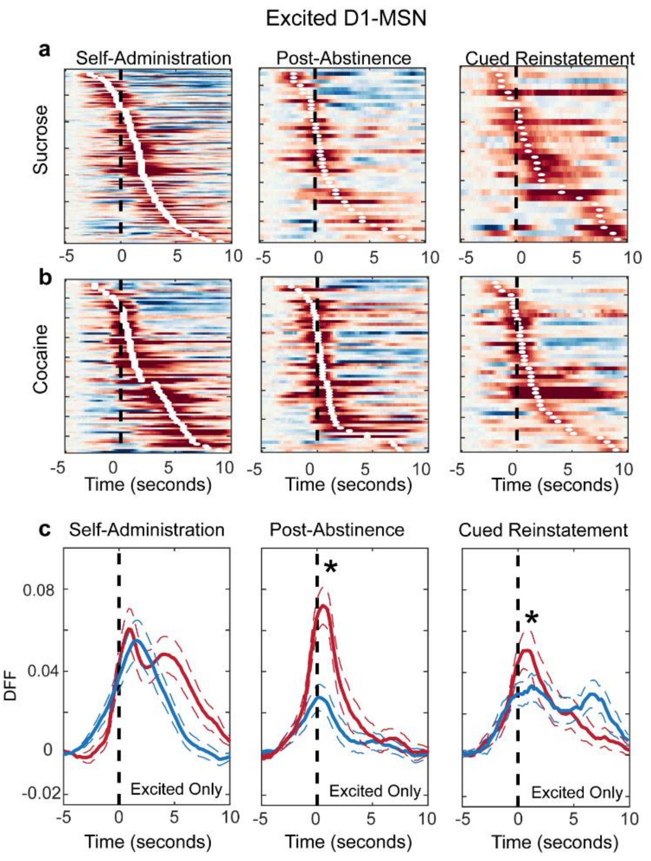


**Figure S5. Excited D1-MSNs subpopulations during cocaine and sucrose self-administration, post-abstinence seeking and cued reinstatement.**

**(a-b)** Heatmaps showing excited D1-MSN during (a) sucrose and (b) cocaine self-administration, post-abstinence seeking and reinstatement. The neurons were sorted according to the location of their peak activity in the recorded window. White dots on every row mark the location of the peak activity of each neuron. **(c)** Mean activity of time-locked excited D1-MSNs during self-administration, post-abstinence seeking and cued reinstatement comparing cocaine (red) to sucrose (blue). Cocaine results in a higher amplitude response of excited D1-MSNs during post-abstinence seeking and cued-reinstatement. *$p<0.05$, comparing cocaine to sucrose by have non-overlapping confidence intervals in their respective shuffled distributions.

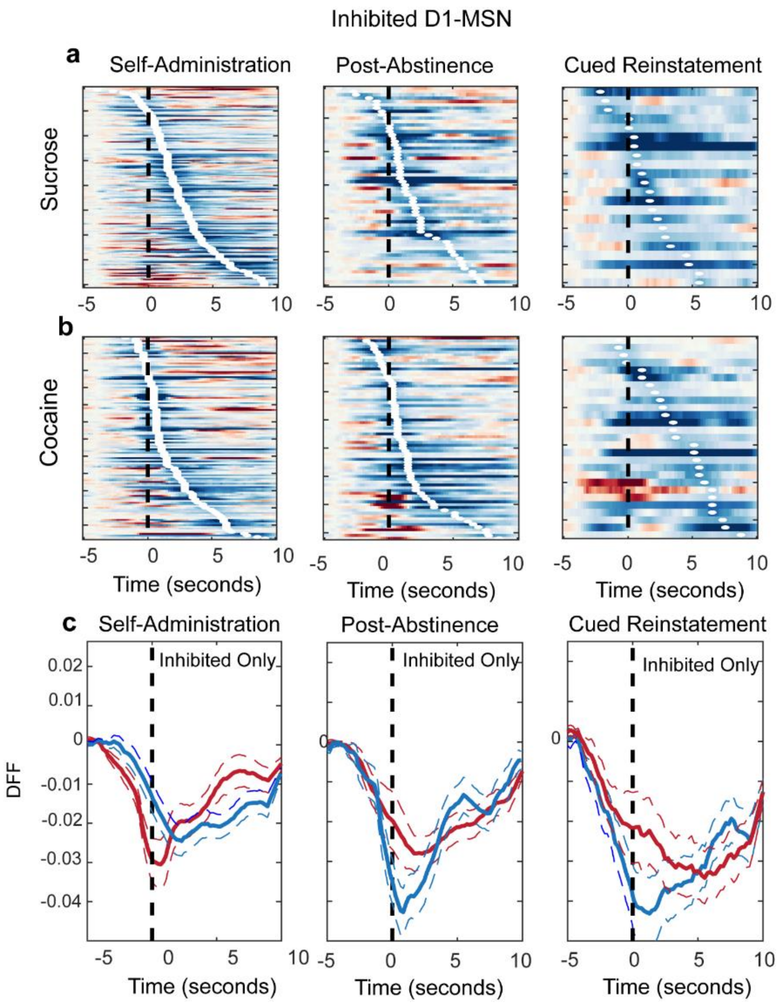


**Figure S6. Inhibited D1-MSNs subpopulations during cocaine and sucrose self-administration, post-abstinence seeking and cued reinstatement.**

**(a-b)** Heatmaps showing inhibited D1-MSN during (a) sucrose and (b) cocaine self-administration, post-abstinence seeking and reinstatement. The neurons were sorted according to the location of their peak activity in the recorded window. White dots on every row mark the location of the lowest activity of each neuron. **(c)** Mean activity of time-locked inhibited D1-MSNs during self-administration, post-abstinence seeking and cued reinstatement comparing cocaine (red) to sucrose (blue). Sucrose results in a more pronounced decrease in activity of inhibited D1-MSNs during post-abstinence seeking and cued-reinstatement.

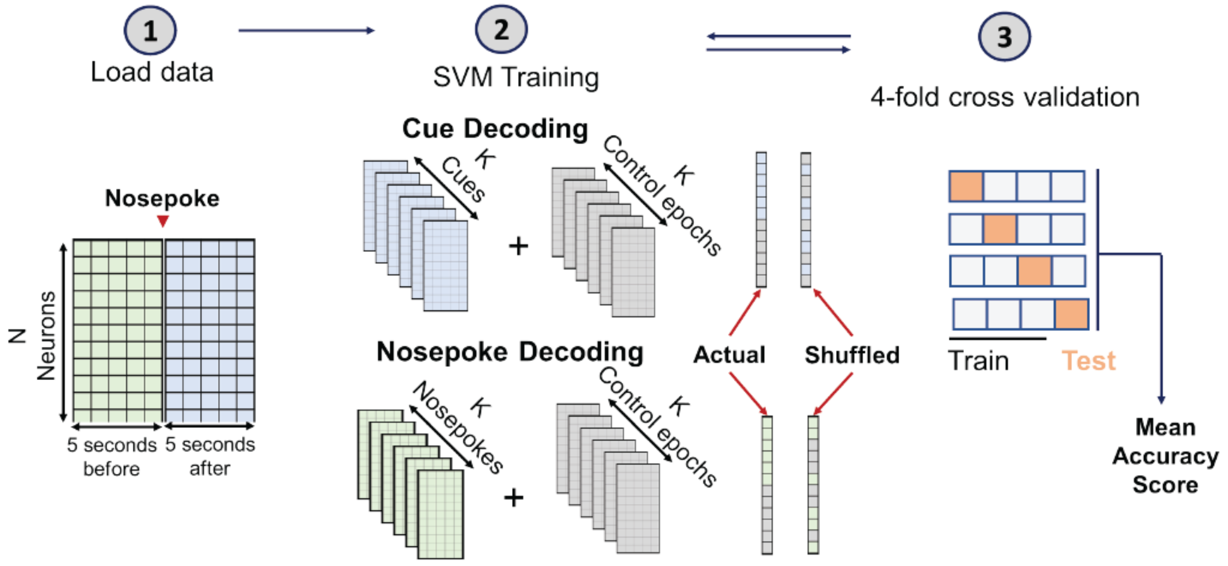


**Figure S7. Illustration of machine learning protocol to decode behavior.** Neuronal data around 5 seconds before and 5 seconds after the nosepoke were extracted and sorted into N-by-F-by-k matrices, where N is the number of neurons, F is the number of frames across 5 seconds, and K is the number of cues received by the animal during the session used. An equal number of control epochs was generated from the data, during random points during the task, and was used as negative control epochs to train the SVM classifier. SVM classifier was trained twice using real labels or shuffled labels of the epochs. SVM training was done using 4-fold cross validation. The difference between the mean accuracy score of the classifier using actual data and shuffled data were reported.
